# PEG-Free Polyion Complex Nanocarriers for Brain Derived Neurotrophic Factor

**DOI:** 10.1101/2022.05.23.492849

**Authors:** James M. Fay, Chaemin Lim, Anna Finkelstein, Elena Batrakova, Alexander V. Kabanov

## Abstract

Many therapeutic formulations incorporate poly(ethylene glycol) (PEG) as a stealth component to minimize early clearance. However, PEG is immunogenic and susceptible to accelerated clearance after multiple administrations. Here, we present two novel reformulations of a polyion complex (PIC), originally comprised of poly(ethylene glycol)_113_-b-poly(glutamic acid)_50_ (PEG-PLE) and brain derived neurotrophic factor (BDNF), termed Nano-BDNF (Nano-BDNF PEG-PLE). We replace the PEG based block copolymer with two new polymers, poly(sarcosine)_127_-b-poly(glutamic acid)_50_ (PSR-PLE) and poly(methyl-2-oxazolines)_38_-*b*-poly(oxazolepropanoic acid)_27_-*b*-(poly(methyl-2-oxazoline)_38_ (PMeOx-PPaOx-PMeOx) which are driven to association with BDNF via electrostatic interactions and hydrogen bonding to form a PIC. Formulation using a microfluidic mixer yields small and narrowly disperse nanoparticles which associate following similar principles. Additionally, we demonstrate that encapsulation does not inhibit access by the receptor kinase, which effects BDNF’s physiologic benefits. Finally, we investigate the formation of nascent nanoparticles through a series of characterization experiments and isothermal titration experiments which show the effects of pH in the context of particle self-assembly. Our findings indicate that thoughtful reformulation of PEG based, therapeutic PICs with non-PEG alternatives can be accomplished without compromising the self-assembly of the PIC.

## 1. Introduction

Nanoformulations incorporating poly(ethylene glycol) (PEG) coronas have enabled immune evasion and the subsequent delivery of packaged therapeutic agents [1,2] However, despite PEG’s status as the “gold standard” stealth polymer, several issues have been identified. Accelerated clearance after repeat administration of stealth nanoparticles due to elicitation of antibodies has long been noted in the literature [3]. Furthermore, possibly due to the ubiquity of PEG in pharmaceutical products and consumer goods, much of the population has existing α-PEG antibodies [4], which could yield accelerated clearance of PEG based pharmaceutical agents or allergic reactions to critical therapies. Recent reports of possible allergic reactions to PEG in the Pfizer and Moderna vaccines against SARS-COVID-2 have brought an already expansive body of work to the forefront of the scientific consciousness [5–9]. PEG’s prevalence in modern therapeutics and immunogenicity can combine to create a population level problem as an increasing number of widely distributed therapies incorporate PEG.

Many groups, our own included, have presented a number of novel nanoformulations whereby small molecules, nucleic acids, and proteins are protected by PEG coronas. In one particular example, we presented a polyion complex (PIC) comprised of poly(ethylene glycol)_113_-b-poly(glutamic acid)_50_ (PEG-PLE) and brain-derived neurotrophic factor (BDNF), termed Nano-BDNF PEG-PLE, as a therapeutic candidate for the treatment of neurodegenerative diseases [10,11]. BDNF has demonstrated great promise in treating animal models of several diseases including ischemic stroke [10], Alzheimer’s disease [12], Parkinson’s disease [11,13], etc. [14]. However, delivery of BDNF to the brain or across the blood-brain barrier (BBB) remains difficult, despite BDNF’s ubiquity in the body. Endogenous trans-BBB transporters are quickly saturated, serum half life is short, and direct administration of BDNF to the brain is attenuated by rapid efflux [11,15]. Nano-BDNF PEG-PLE nanoparticles effectively mediated the delivery of BDNF to its target receptor after intranasal to brain administration [11]. Additionally, we have demonstrated that Nano-BDNF PEG-PLE presents clear benefits over unformulated BDNF in the treatment of Parkinson’s disease and ischemic stroke models in mice while preserving BDNF’s circulation and residence in affected areas [10,11].

PEG is a long, flexible, and hydrophilic polymer which can comprise a corona which entropically exclude opsonins from penetrating into the nanoparticle and interacting with the therapeutic cargo [16–18]. Similar polymers have shown the same stealth effect allowing nanoparticles to circulate longer and minimizing unwanted clearance or surface fouling [19–21]. Work from our own lab and others has identified a variety of polymers which recapitulate the long, hydrophilic, and flexible nature of PEG [19,22–27]. In particular, our group has extensive experience utilizing poly(2-oxazolines) which present a useful platform for the production of functionalized block copolymers [19,28–34]. Convenient extension of polymers using living-cationic-ring-opening-polymerization (LCROP) enables precise control of block length and organization. Several advancements using polyoxazoline constructs to form drug carrying nanoparticles which demonstrate a stealth effect similar to PEGylated liposomes or PICs have been presented [19,24,37,29–36]. Similarly, other groups have used poly(sarcosine) (PSR), another flexible and hydrophilic polymer, to similar effect in non-fouling surfaces and stealth nanoparticles [20,26,38-42].

Here we introduce two types of PIC’s similar to the original Nano-BDNF PEG-PLE but free of PEG. A nanoparticle comprised of BDNF and poly(sarcosine)_127_-b-poly(glutamic acid)_50_ (Nano-BDNF PSR-PLE) and a series of nanoparticles comprised of BDNF and poly(methyl-2-oxazolines)_x_-*b*-poly(oxazolepropanoic acid)_y_-*b*-(poly(methyl-2-oxazoline)_z_ (PMeOx_x_-PPaOx_y_-PMeOx_z_) (B1-5) (Nano-BDNF B1-5) are presented. We examine the formulation of several Nano-BDNF species and present two nanoparticles which are small and narrowly dispersed. We also pursue a deeper understanding of the thermodynamics of association between BDNF and our various polymers as well as their behavior to demonstrate similarity between our three chosen polymers. Finally, we consider the application of the nanoparticles to cells stably transfected with tropomyosin receptor kinase B (TrkB), the primary receptor for BDNF [43], in order to assure that we do not prohibit activation of the desired target. In conclusion, we present two novel non-PEG formulations which demonstrate similarity to the original Nano-BDNF PEG-PLE formulation in morphology, activity, and thermodynamic behavior with potential for therapeutic effect in neurological diseases.

## 2. Materials and Methods

### 2.1 Materials

Methyl triflate, 2-methyl-2-oxazoline, methoxycarboxyethyl-2-oxazoline, and piperidine were obtained from Sigma Aldrich. Dialysis membrane with molecular weight cut off [MWCO] 3,500 was purchased from Spectra/Pore. The following materials were purchased from Thermo/Fisher: WhatMan Anotop 0.025 μm non-sterile filters, 25 gauge 0.5 inch, long blunt needle (Strategic Applications inc.), 40% acrylamide 29:1, TEMED, Ammonium persulphate, Malvern Folded Cap Cells, DLS cuvettes (Malvern), radioimmunoprecipitation assay buffer, HALT protease and phosphatase inhibitor cocktail, SuperSignal™ West Pico PLUS Chemiluminescent Substrate, iBRIGHT prestained protein ladder, Geneticin™ Selective Antibiotic (G418 Sulfate), Neurobasal Media, B27 supplement, Glutamax 100X, acetonitrile, potassium carbonate, diethyl ether, methanol, agarose tablets, and sodium hydroxide. BDNF was obtained from Novoprotein which later became Bon Opus Biosciences. PSR-PLE was obtained from Peptide Solutions LLC. Colorado calf serum was obtained from the Colorado Serum Company. We obtained Bio-Safe Coomassie Stain and Native Sample buffer for protein gels 4X from Bio-Rad.

### 2.2 Polymer Synthesis

Under dry and inert conditions achieved in MBraun glovebox with N2 atmosphere, 1 molar equivalent of the polymerization initiator, methyl triflate (MeOTf), and 10 or 30 molar equivalents of 2-methyl-2-oxazoline (MeOx) were dissolved in 2 mL of acetonitrile (ACN). The mixture was sealed and moved to react in a microwave for 15 min at 150 W and 130 °C. After cooling, the solution was returned to dry and inert conditions where 10 or 30 molar equivalents of the monomer for the second block, methoxycarboxyethyl-2-oxazoline (MestOx), were added. The mixture was sealed and moved to the microwave following the same procedure as above. After cooling, the procedure was repeated with 10 or 30 molar equivalents of MeOx. Polymerization was terminated by adding 3 molar equivalents of piperidine and microwaving for 1 hour at 150 W and 40°C. An excess of K2CO3 was added to ensure complete cessation of the reaction. The solution was stirred overnight, filtered through a 0.45 μm filter, precipitated in diethyl ether, collected, and freeze dried. The product was dissolved in methanol and 0.1 N NaOH was added to 1 molar equivalent NaOH with the methyl ester group on the poly(MestOx) block of the copolymer to convert to PPaOx. The resultant solution was allowed to react for 2 hours at 80°C. The solution was then dialyzed against distilled water using a dialysis membrane (Spectra/Pore; molecular weight cut off [MWCO] 3,500), product was freeze dried for storage until use.

### 2.3 Polymer Characterization by ^1^H NMR

Copolymers were suspended in deuterated methanol (MeOD) and ^1^H nuclear magnetic resonance (NMR) spectra were obtained using an INOVA 400 NMR machine. Data were analyzed using MestReNova (11.0). Spectra were calibrated using the MeOD solvent signal at 4.78 ppm. The number average molecular weight was determined by calculating the ratio of the initiator and each repeating unit.

### 2.4 Preparation of PIC by Manual Mixing

The formulation was manufactured as described previously in Jiang et al.^6^ Briefly, a concentrated solution of a block copolymer (0.1, 1, or 10 mg/mL) in a specified buffer was added dropwise to the BDNF solution ([BDNF] =1 mg/mL); each drop interspersed with a gentle vortex for 10 seconds. The solution was then diluted to the desired concentration using the buffer and vortexed for another 30 seconds, after which the solution was incubated for 30 min at room temperature.

### 2.5 Preparation of PIC Using a Microfluidic Mixer

Separate solutions of BDNF and copolymer in a specified buffer were prepared at 2X the desired end concentrations. We used either phosphate buffered saline (PBS) (10 mM NaH_2_HPO_4_, 1.8 mM KH_2_PO_4_, 2/7 mM KCl, 137 mM NaCl, pH 7.4)) or lactate buffered Ringer’s solution (LR) (100 mM NaCl, 3.7 mM KCl, 1.3 mM CaCl_2_, 1.2 mM MgCl_2_, 28 mM NaHCO_3_, 20 mM NaC_3_H_5_O_3_, pH 7.4). The BDNF and copolymer solutions were loaded into 1 mL syringes which were then connected to the microfluidic mixer apparatus (**Fig. S1**) described previously [11]. The solutions were each injected into the mixer at 50 μL/min to mix at a 1/1 volume ratio. The first 100 μL were discarded to allow only properly mixed samples to be collected before experiments.

### 2.6 Nanoparticle Size, Dispersity, and Zeta Potential

Nano-BDNF was prepared at the desired Z_-/+_ ratio. The particle size and polydispersity index (PDI) were analyzed by dynamic light scattering (DLS) using the Zetasizer Nano ZS (Malvern Instruments Ltd.) with a scattering angle of 173° at RT. For all DLS measurements, the BDNF concentration was 0.05 mg/mL. Unless specified otherwise, the DLS sizes are reported as intensity weighted (z-averaged) diameters. The nanoparticle tracking analysis (NTA) was performed using the NanoSight NS500 (Malvern Panalytical) or Zetaview (ParticleMetrix). All samples were prepared at BDNF concentration 0.005 mg/mL. For the measurements of particle size and distribution using the NanoSight NS500, the samples were stored on ice and allowed to return to RT prior to use. Readings were taken under flow for 60 s at 20 μL/s using the syringe pump. For the measurements of particle size, distribution, and zeta potential using the Zetaview, the samples were prepared at BDNF concentration 0.005 mg/mL and analyzed either immediately or incubated at RT for various times. Before the measurements, these samples were diluted 10X. Both NanoSight NS500 and Zetaview are reported as number-weighted diameters.

### 2.7 Horizontal Agarose Gel Electrophoresis (HAGE)

Agarose gels (0.5 % wt.) were prepared using 20 mM MOPS buffer, containing 2 mM sodium acetate, pH 7.4 which was also used as the running buffer. Nano-BDNF was manually prepared as described above in LR solution, at a concentration of 0.075 mg/mL and volume of 18 μL. The sample was diluted 2X with Native Loading Buffer, pH6.8 (Bio-Rad) and 35 μL of the final solution were loaded into the wells. Electrophoresis was performed with a constant voltage of 80 V for 1 hour. The gel was washed in DI water 2X and stained using Biosafe Coomassie Stain (Bio-Rad) for at least 1 hour. Destain was performed using DI water until the gel was mostly clear. Images were taken using the FluorChem E machine (ProteinSimple).

### 2.8 Isothermal Titration Calorimetry (ITC)

BDNF and polymer solutions were dialyzed against 10 mM phosphate, pH 7.4, 10 mM phosphate, pH 5.8, or 10 mM sodium acetate, pH 5.0 overnight with at least two buffer changes. BDNF was diluted to 10 μM and the polymers were diluted to the indicated concentrations then stored on ice. Samples were loaded into the MicroCal PEAQ-ITC machine (Malvern Panalytical) and stored at 4°C in the tray. The cell and syringe were kept at 25°C and the experiment was performed at 25°C. The polymer was added in 2 μL steps except for the first addition which was 0.2 μL. The cell was allowed to incubate for 3 minutes between each step. Data analysis was performed using the MicroCal ITC analysis software in Origin.

### 2.9 Potentiometric Titration of Polymers

Polymers were solubilized in 10 mL of nitrogen flushed dH2O and a 1.1 molar equivalent aliquot of 1.1 M HCl was added to the 10 mL of polymer solution at a concentration of 500 μM ionizable units. 50 mM NaOH was added in 10 μL increments, the pH was allowed to stabilize (~1 min dwell time), and readings were taken using an Oaklon ion 700 pH meter with Fisherbrand™ accumet™ glass electrode.

### 2.10 Cell culture

NIH 3T3 cells stably transfected with TrkB (NIH 3T3 TrkB) were cultured in GlutaMax DMEM supplemented with 10% calf serum, G418 (100 μg/mL), penicillin (100 μg/mL), and streptomycin (100 μg/mL). Cells were cultured at 37°C in a humidified CO_2_ (5%) incubator.

### 2.11 BDNF Stimulation

The NIH 3T3 TrkB cells were plated at 5×10^5^ cells/well on 6-well plates 24 hours before treatment. Five hundred nanograms of BDNF or equivalent amount of Nano-BDNF was added to the wells then incubated for 5 min at 37°C in the incubator. Formulations were prepared via microfluidic mixing at Z_-/+_ = 5 in cell culture media. Media was removed and replaced with radioimmunoprecipitation assay buffer with HALT™ proteinase and phosphatase inhibitors (ThermoFisher), then the plate was moved to 4°C and agitated using a plate shaker. Material was collected and separated into supernatant and pellet. The supernatant was measured for protein concentration by the BCA assay, then 25 μg of protein was loaded into each well and electrophoresed through a 12.5% SDS-PAGE gel, transferred to PVDF membrane, and visualized via horseradish peroxidase (Supersignal™ West Pico Plus Chemiluminescent Substrate) following standard western blotting procedures. The following antibodies were used to visualize their cognate proteins: Phospho-p44/42 MAPK (Erk1/2) (Thr202/Tyr204) Antibody (CST 9101), p44/42 MAPK (Erk1/2) Antibody (CST 9102), Anti-Rabbit HRP (CST 7074S).

## 3. Results

### 3.1. Manufacture and Characterization of Poly(2-oxazoline) Block Copolymers

Previous work demonstrated that BDNF readily associated with the negatively charged PEG-PLE diblock copolymer to form stable nanoparticles [10,11]. Pursuant to our objective of replacing PEG with alternative polymers, we produced four block copolymers using LCROP as described previously [44] (**Fig. 1**). The new polymers were designed as anionic A-B-A type block copolymers composed of hydrophilic uncharged PMeOx (A block) and negatively charged PPaOx (B block). The amounts of monomers, ten or thirty in each step, added during synthesis were selected to produce four variations of PMeOx-PPaOx-PMeOx with the following patterns: short-short-short, short-long-short, long-short-long, and long-long-long (**Fig. 2**). Structure and length ratio were confirmed via ^1^H-NMR (**Fig. 1,** supplementary **Fig. S2**). As an alternative approach to formulating BDNF, we used an anionic PSR-PLE di-block copolymer that was available commercially with a similar structure to the original PEG-PLE polymer used in our previous study (**Fig. 2**).

**Figure 1.**
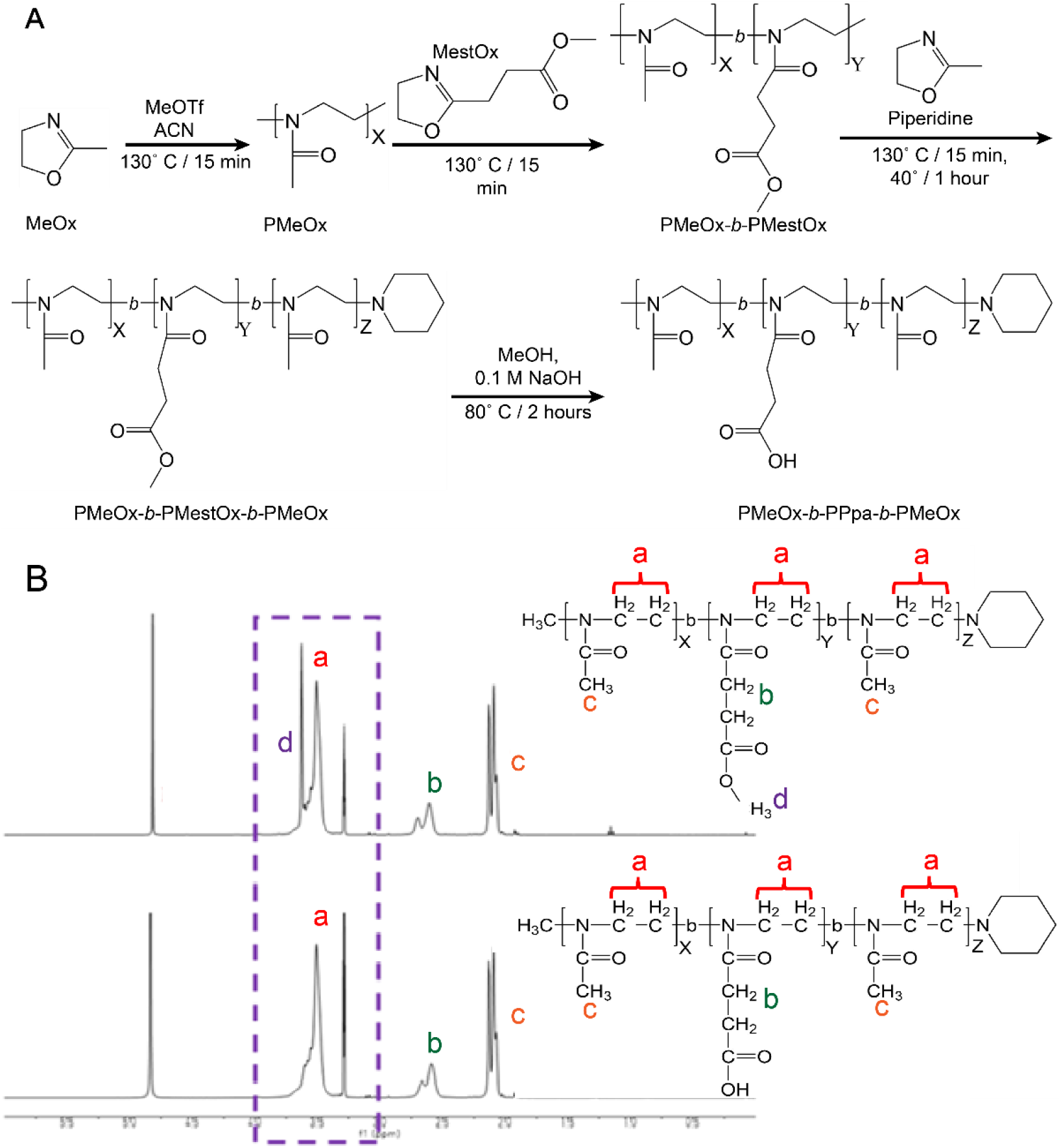
Polymer synthesis and characterization. **(A)** Synthesis scheme for the preparation of poly(2-oxazoline) tri-block copolymers via LCROP. Briefly, the initiator, MeOTf, was reacted with MeOx in ACN for 15 min at 130°C followed by sequential polymerization of subsequent blocks in ACN for 15 min each then termination with piperidine reacted for 1 hour at 40°C. The protective methyl group on the MestOx block was removed by reacting with NaOH in MeOH for 2 hours at 80°C. **(B)** ^1^H-NMR peak assignments of PMeOx-*b*-MestOx-*b*-PMeOx and PMeOx-*b*-PPaOx-*b*-PMeOx. Note the elimination of peak **(d)** representing the deprotection of the hydroxyl group on the middle block. Spectra of the four polymers produced can be found in the supplement (**Fig. S2**).

**Figure 2.**
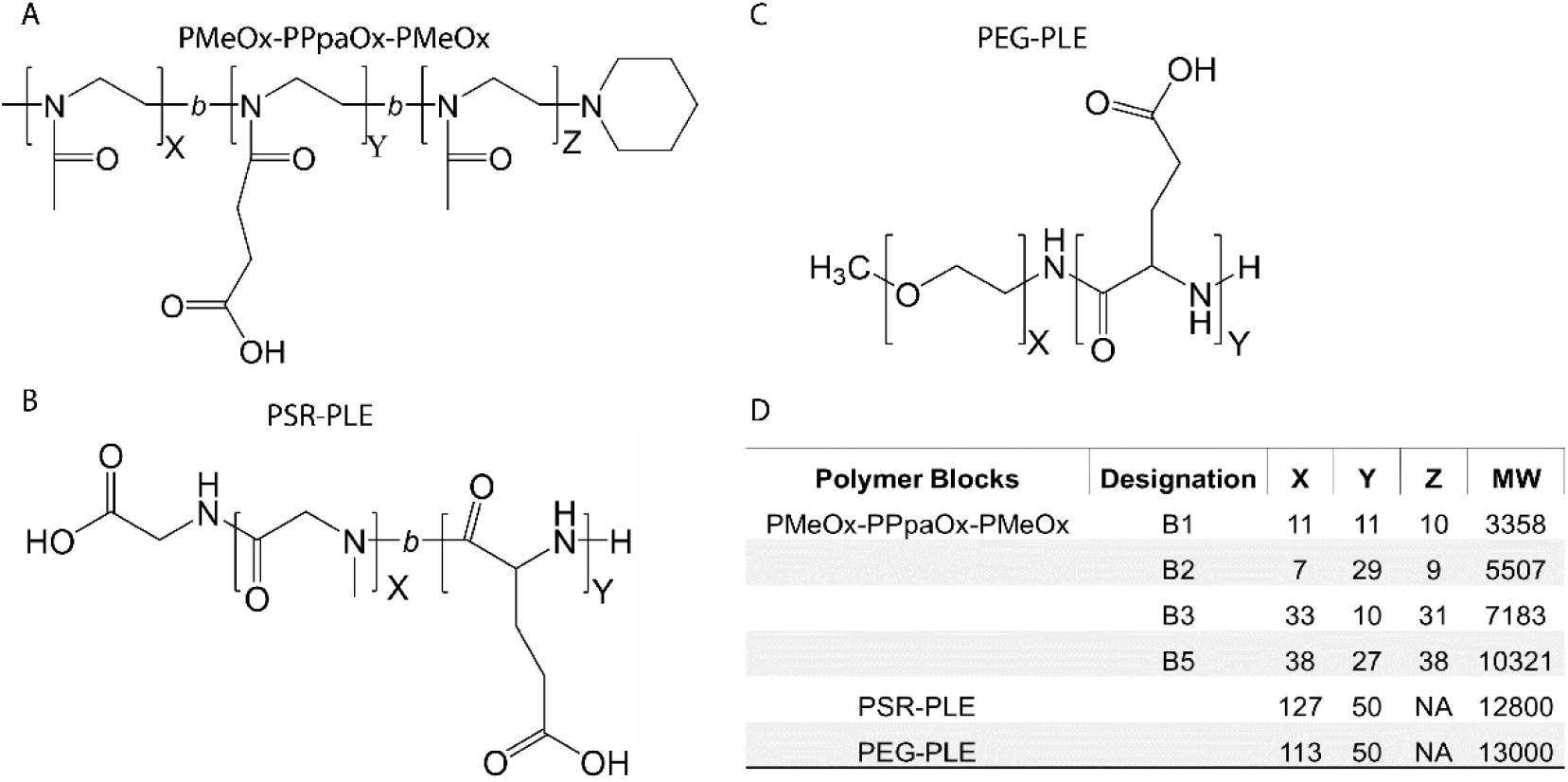
Block copolymer structures **(A-C)**, and molecular characteristics **(D)**.

### 3.2. Nano-BDNF Candidate innowing

The length and structure of the copolymer blocks can drastically affect self-assembly of block copolymers with proteins or other cargo for delivery [45]. In addition, the mode of mixing is of paramount importance for formulation of PIC that can form non-equilibrium, slowly transitioning aggregates [22,46]. This is characteristic of PIC’s that contain proteins or peptides, such as BDNF, that can self-aggregate because of intermolecular hydrophobic interactions or hydrogen bonding. To quickly identify which block copolymers are most suitable for formulation, we adopted a winnowing strategy. Towards that end, we formulated Nano-BDNF in 10 mM Hepes, pH 7.4 to interrogate block copolymer-protein interaction and identify which copolymers yield relatively small, narrowly dispersed particles.

The ionic strength, salt identity, and pH are all expected to strongly affect the formation of the PIC particles. Initial formulation in minimal conditions of only HEPES buffer allows us to identify which copolymers form stable PICs with BDNF in the near absence of salts. We initiated by manually formulating Nano-BDNF in 10 mM HEPES, pH 7.4, by mixing each copolymer and BDNF. In addition to the effects of buffer components, the ratios in which copolymer and protein are mixed may strongly affect the formation of particles in terms of size, dispersity, stability, etc.^6^ The mixing ratio between carboxyl groups of copolymer and amino groups of BDNF is defined here as follows:

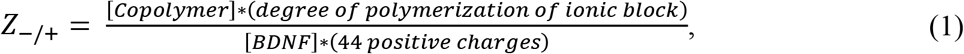

Each PIC was formulated at several Z_-/+_ ratios (1, 2, 5, 10). We assessed the size and dispersity of the particles using DLS over two days. The poly(2-oxazoline) tri-block copolymers showed major differences in their propensity to associate with BDNF. While we observed particle formation in all cases, copolymers B2 and B5 yielded smaller and more uniform particle populations (**Fig. 3** and supplementary **Fig. S3-S12**). Formulation with B1 formed large aggregates and B3 produced highly disperse particles. Therefore, we disregarded those copolymers in future studies and focused on B2 and B5. We noted that the formulations containing B2 and B5 (at several Z_-/+_ ratios 2, 5, 10) were variable and poorly reproducible, indicating the capricious nature of the manual mixing. Nano-BDNF formulated with PSR-PLE was also not greatly amenable to the formulation at this initial stage as mixing formed highly disperse PIC particles (**Fig. 3,** supplementary **Fig. S3**). However, we continued optimization using PSR-PLE due to similarity to the structure of PEG-PLE used in Jiang, et al [11].

**Figure 3.**
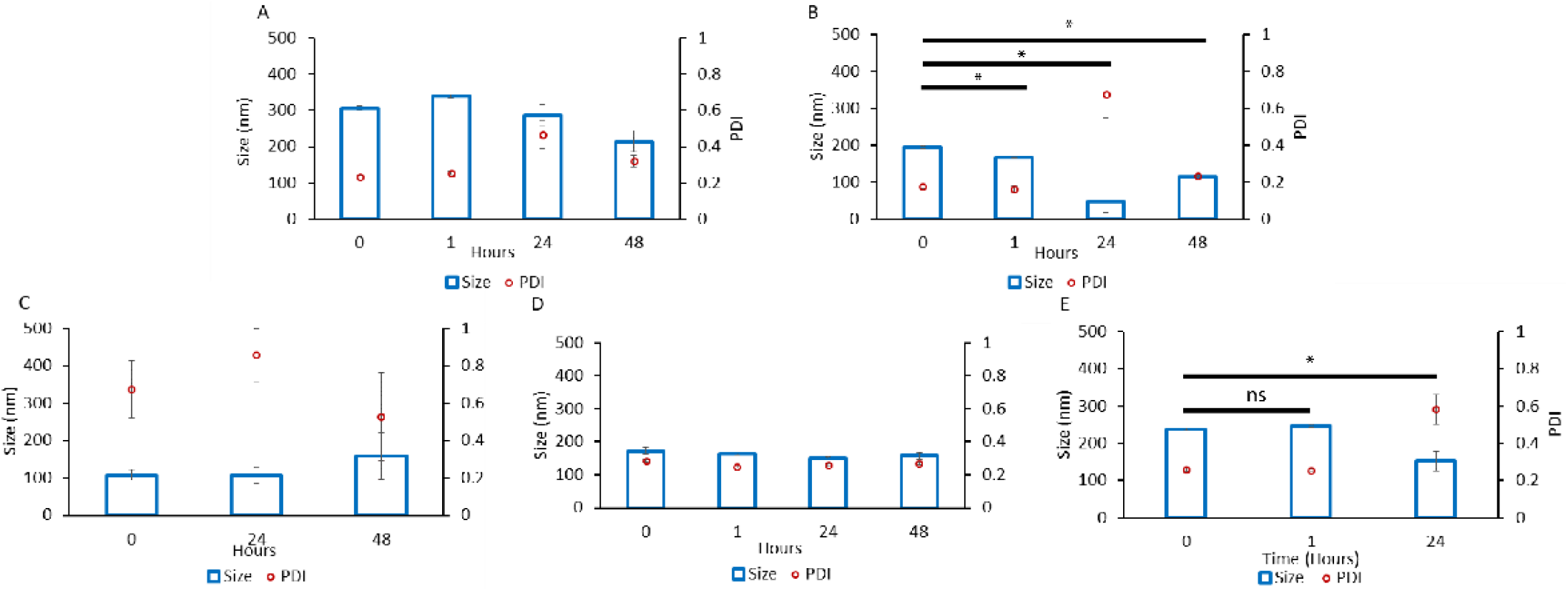
Particle size and dispersity of Nano-BDNF. The complexes were prepared by manual mixing of the solutions of BDNF and **(A)** B1, **(B)** B2, **(C)** B3, **(D)** B5, or **(E)** PSR-PLE in 10 mM HEPES, pH 7.4. The component charge ratio was Z_-/+_ = 1 **(B-E)** or Z_-/+_ = 2 **(A)**. The DLS measurements were carried out for up to 48 hours. Values are mean ± SEM, * p < 0.05, unmarked plots do not show statistically significant differences. Statistical significance was established using an unpaired two tailed Student’s t-test.

The effects of low molecular-mass electrolytes on PICs have been noted in the literature such that different salts can affect particle formation, size, and dispersity [45]. To assess these effects in our systems we opted for the simple addition of 150 mM NaCl to a 10 mM HEPES, pH 7.4 solution. Nano-BDNF was prepared using B2, B5, and PSR-PLE via manual mixing. We did not observe a decisive advantage using any one copolymer under these conditions, B2 yielded small particles with very high polydispersity, PSR-PLE yielded very large but narrowly disperse particles, and B5 was a mix (**Fig. 4**). Most notably, the particles formed at higher Z-/+ ratios using B5 and PSR-PLE (**Fig. S13 E, F, I**) were smaller and less dispersed than in the absence of added salt. It may be that the addition of NaCl allowed ion exchange and facilitated packing. The number weighted particle size for B2 was negligible compared to the intensity weighted particle size indicating only small number of aggregates existed that in the presence of salt (**Fig. S14 A, B, C**).

**Figure 4.**
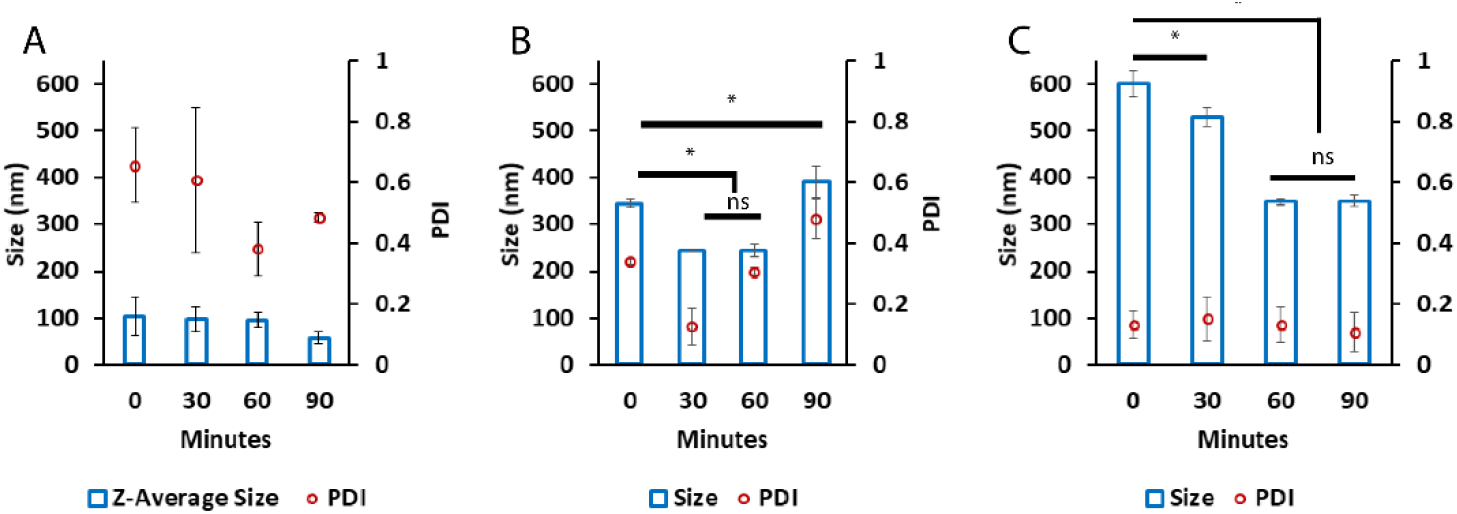
Particle size and dispersity of Nano-BDNF in the presence of small-molecular mass electrolyte. All Nano-BDNF samples were prepared at Z_±_ = 5 by manual mixing of the solutions of BDNF and **(A)** B2, **(B)** B5, or **(C)** PSR-PLE in 10 mM HEPES, pH 7.4, 150 mM NaCl. The DLS measurements were carried out for over 2 hours. Values are mean ± SEM, * p < 0.05, unmarked plots do not show statistically significant differences. Statistical significance was established using an unpaired two tailed Student’s t-test.

To select the best copolymers to continue optimization, we altered formulation methodology to include microfluidic mixing, which has yielded decreased particle size and dispersity in past publications [11,47,48]. B2, B5, and PSR-PLE were incorporated into Nano-BDNF using microfluidic mixing in 10 mM HEPES, pH 7.4 with no elementary salt added. We measured size and dispersity using DLS and NTA (**Fig. 5**). Using microfluidic mixing we generally observed smaller and less disperse particles (**Fig. 5**) when compared to manual formulation in the same buffer (**Fig. 3**, and supplementary **Fig. S3-S12**). PSR-PLE particles formed using this technique considerably decreased the size and narrowed dispersity compared to manual mixing (**Fig. 3**). This result is consistent with previous work where Nano-BDNF formulated with PEG-PLE yielded smaller, more narrowly disperse particles when formulated using the same microfluidic mixer. Considering the totality of our results (**Fig. 3–5** and supplementary **Fig. S3-S14**) we elected to pursue formulation with PSR-PLE and B5.

**Figure 5.**
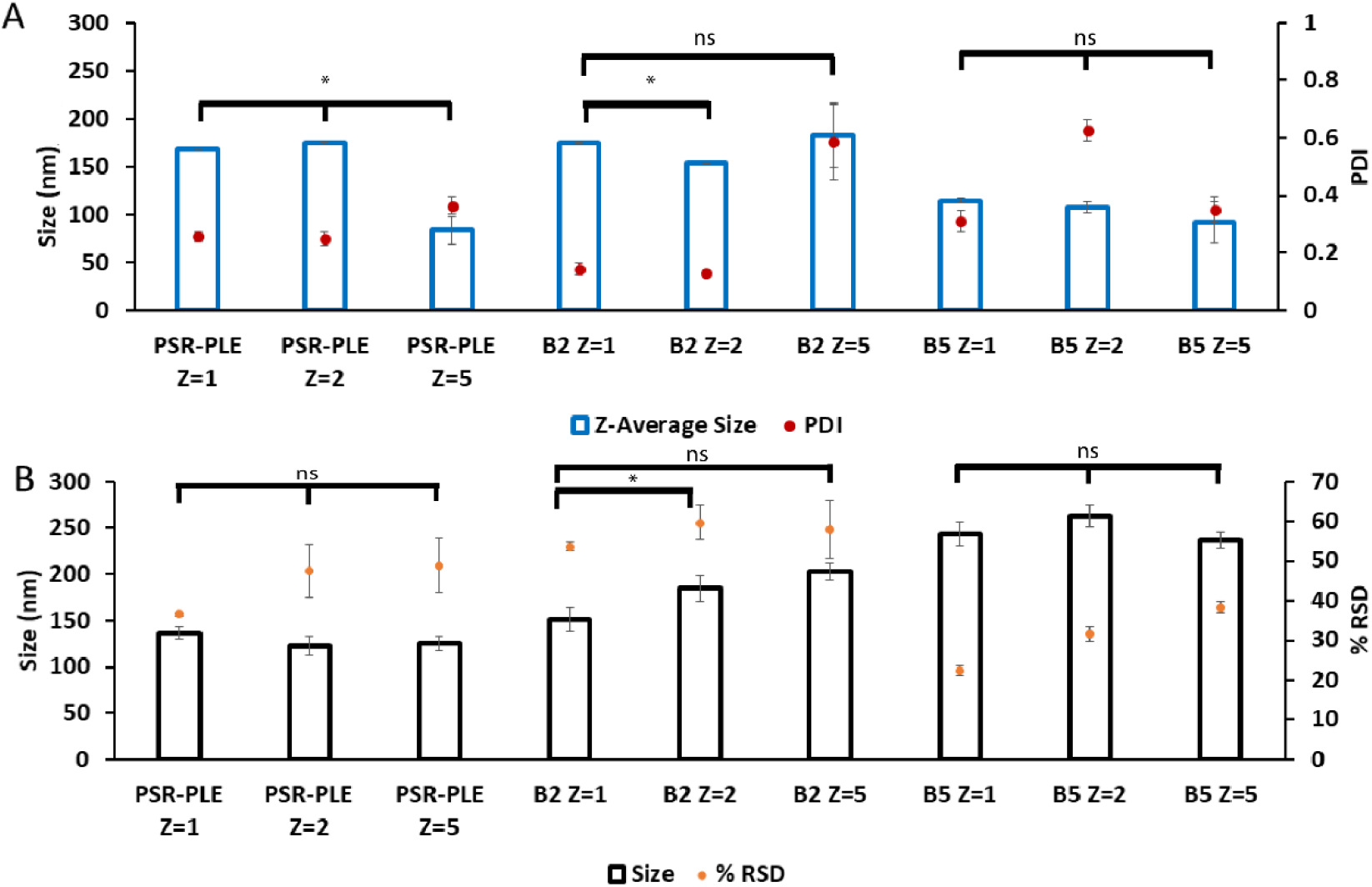
Particle size and dispersity of Nano-BDNF prepared by microfluidic mixing. Nano-BDNF samples were prepared in 10 mM Hepes, pH 7.4 at various Z_-/+_ ratios. The complexes were characterized by **(A)** DLS or **(B)** NTA using Nanosight NS500. Values are mean ± SEM, * p < 0.05, unmarked plots do not show statistically significant differences. Statistical significance was established using an unpaired two tailed Student’s t-test.

### 3.3. Preparation of Nano-BDNF Particles Suitable for Injection

The final stage of optimization was to prepare Nano-BDNF in isotonic solutions formulated with saline that are commonly used for in vivo injections. We selected PBS and LR for formulation and prepared Nano-BDNF in these buffers using copolymers B5 and PSR-PLE via microfluidic mixing. We analyzed the resultant particles size and distribution right after preparation (**Fig. 6**) and over time (**Fig. S15**). Right after mixing, B5 formed the smallest, least disperse particle population when formulated in PBS at Z_-/+_ = 5. However, the formulations in the LR were still promising since Nano-BDNF B5 at Z_-/+_ = 2 and Nano-BDNF PSR-PLE at Z_-/+_ = 5 both yielded particles with sizes less than 120 nm and with RSD under 30. When size and dispersity were measured over time, we found the LR solution yielded smaller sizes and narrower dispersity for both Nano-BDNF B5 and Nano-BDNF PSR-PLE over several hours using (**Fig. S15**). Therefore, we used LR buffer for the next experiments.

**Figure 6.**
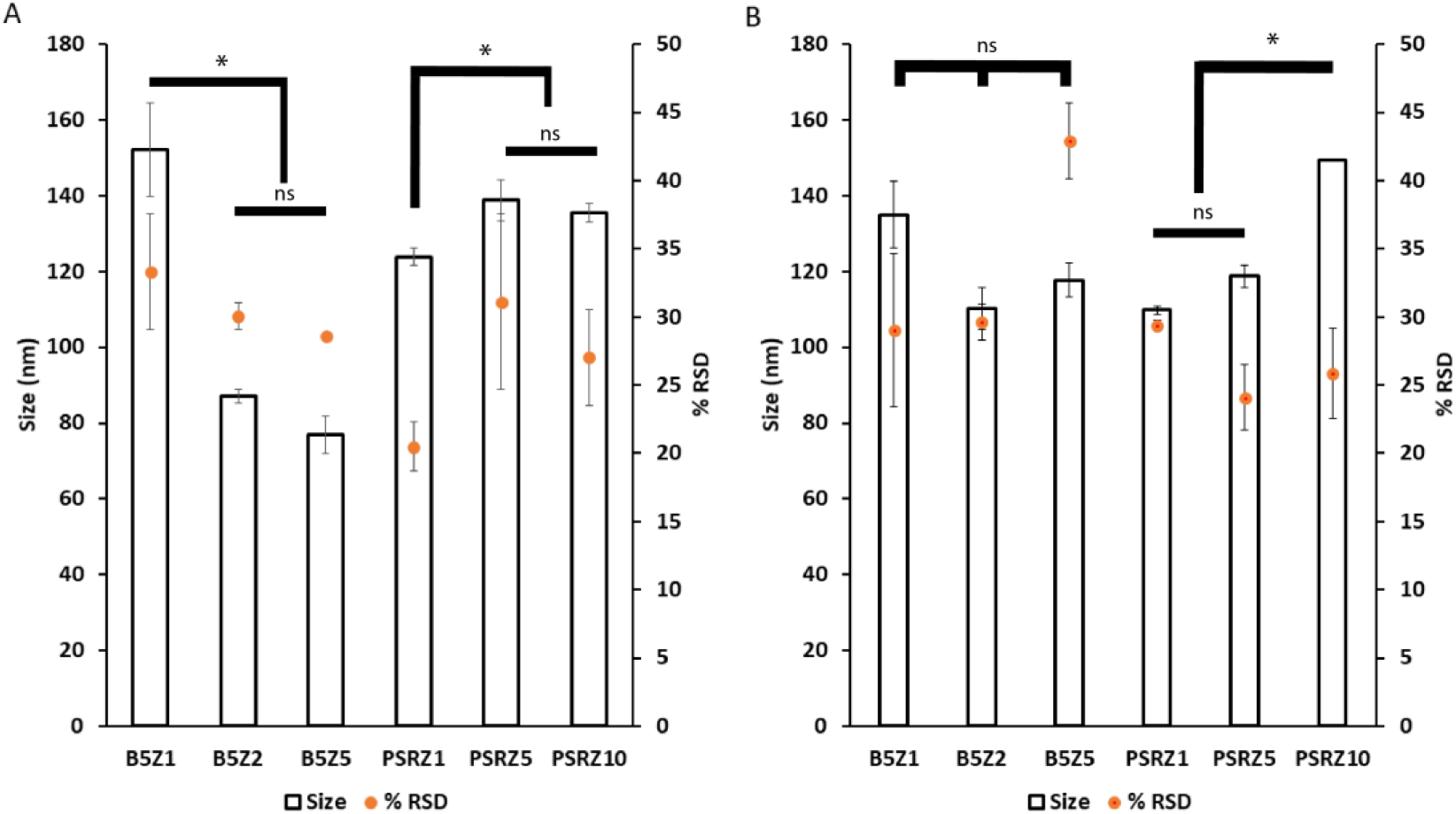
Particle size and distribution of Nano-BDNF samples in isotonic solutions. The complexes were prepared by microfluidic mixing in **(A)** PBS and **(B)** LR. The measurements were carried out within 10 min after mixing at RT. The NTA measurements were carried out using Nanosight NS500. Values are mean ± SEM, * p < 0.05, unmarked plots do not show statistically significant differences. Statistical significance was established using an unpaired two tailed Student’s t-test.

### 3.4. HAGE Visualization of Nano-BDNF

After optimizing the formulation of Nano-BDNF PSR-PLE and Nano-BDNF B5, we visualized their formation and migration through an agarose gel matrix. Jiang, et al. [11] demonstrated that Nano-BDNF PEG-PLE, formulated at excess of polyanion, yielded complexes with a negative overcharge that migrated BDNF towards the cathode despite the protein’s intrinsic net positive charge. Nano-BDNF formulated with PSR-PLE and B5 behaved similarly, such that little or no migration was observed at Z_-/+_ < 5, whereas higher Z_-/+_ (Z_-/+_ = 5, 10) formulations yielded distinct bands migrating towards cathode (**Fig. 7**). Notably, as in Jiang, et al. [11] the free BDNF or Nano-BDNF complexes at Z_-/+_ < 1 remained in the wells with little apparent movement towards anode. We believe that the HAGE approach is unable to distinguish the free protein and complexes formed at deficiency of the copolymer because it is known that BDNF can bind to agarose and is also prone to aggregation [49]. In the complexes formed at lower mixing ratio, the BDNF molecule may not be completely masked by the bound copolymer and partially exposed. On the other hand, the mobility of the complexes formed at higher Z_-/+_ may indicate not only the overcharging but also complete masking of the BDNF by the copolymer that inhibits the aggregation and/or binding to agarose.

**Figure 7.**
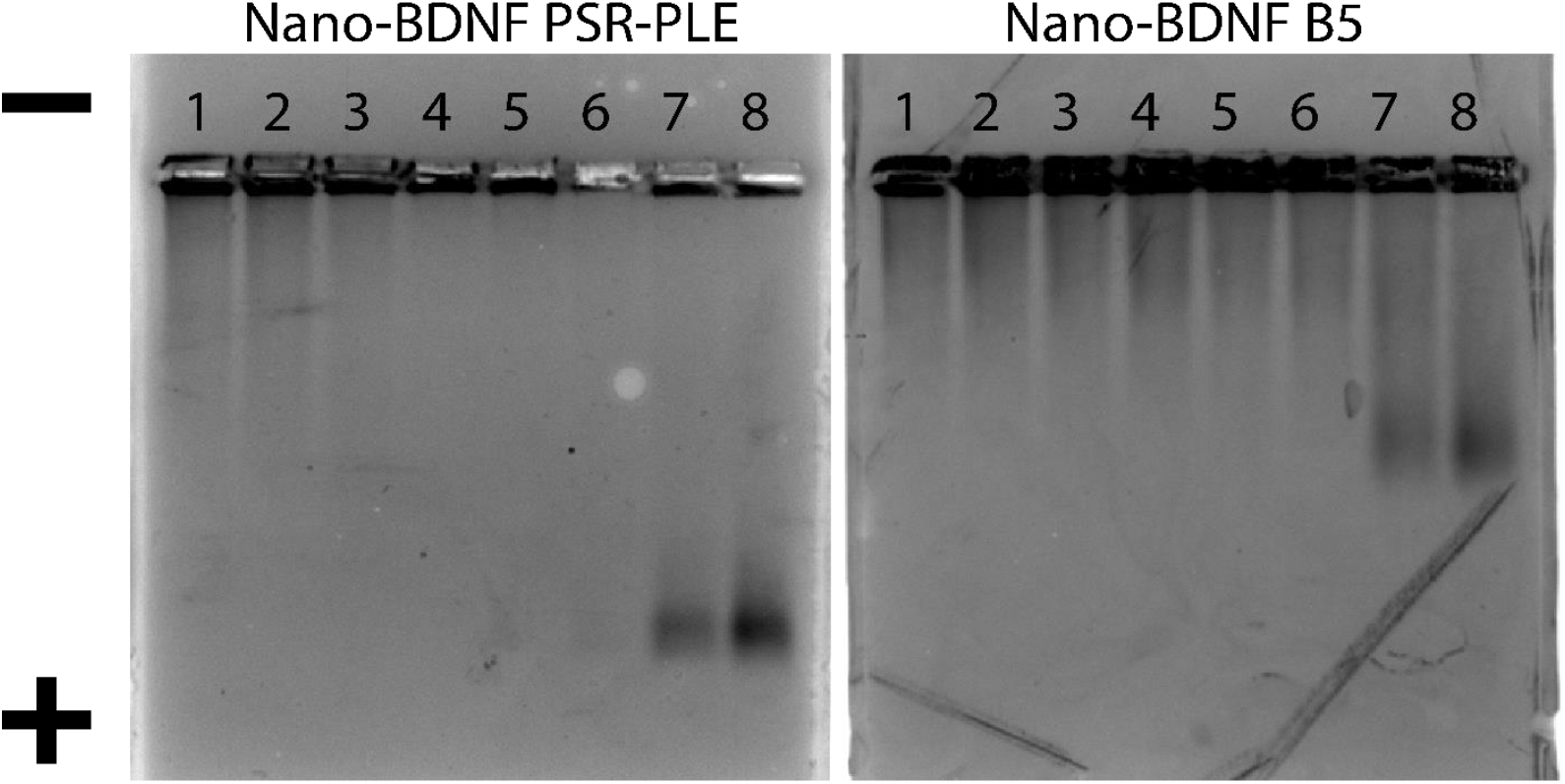
HAGE analysis of Nano-BDNF. The complexes were prepared in LR by manual mixing of BDNF with respective copolymers. Lanes 1 through 8 correspond to BDNF alone (lane 1), and Nano-BDNF prepared at various Z_-/+_: 0.1 (lane 2), 0.25 (lane 3), 0.50 (lane 4), 0.75 (lane 5), 1.0 (lane 6), 5.0 (lane 7), 10.0 (lane 8). The protein bands were visualized by Coomassie blue staining.

### 3.5 Zeta-Potential Characterization of Nano-BDNF

As an unambiguous method for characterization of Nano-BDNF complexes at both low and high mixing ratios, we used the NTA Zetaview machine which directly measures the particle size and zeta potential in the same samples, and additionally reports the calculated dispersity as well as concentration of the particles (**Fig. 8**). For this study we used three block copolymers, PEG-PLE, PSR-PLE and B5, and prepared the PICs by the microfluidic mixing. The overall pattern of behavior of the complexes in 10 mM sodium phosphate, pH 7.4 was very similar for all three copolymers. We were able to observe the formation of the nanosized complexes at Z_-/+_ as low as 0.25 (**Fig. 8A, B**). These complexes were already net negatively charged at low Z_-/+_ ratio but increased in the absolute value of the charge with further addition of anionic copolymer (**Fig. 8C**). For all three copolymers, as the Z_-/+_ increased above 0.5, the concentration of the particles sharply decreased by over 10 times (with no precipitation observed) (**Fig. 8D**). This phenomenon could be explained by the increased binding of the copolymer to BDNF upon increase of Z_-/+_, which promotes self-assembly of PICs with higher molecular masses and higher aggregation numbers of BDNF and copolymer. In contrast, at low Z_-/+_ we may be observing formation of nascent complexes which have lower molecular masses and aggregation numbers. This assumption is only partially supported by the particle size measurements (**Fig. 8A**). While there was a distinct trend for increases in the sizes of Nano-BDNF PEG-PLE and B5, no increases in the particle size were apparent in the case of PSR-PLE. However, the discussed transitions in the PICs may be masked by the shape effects which are not accounted for by our size measurement technique.

**Figure 8.**
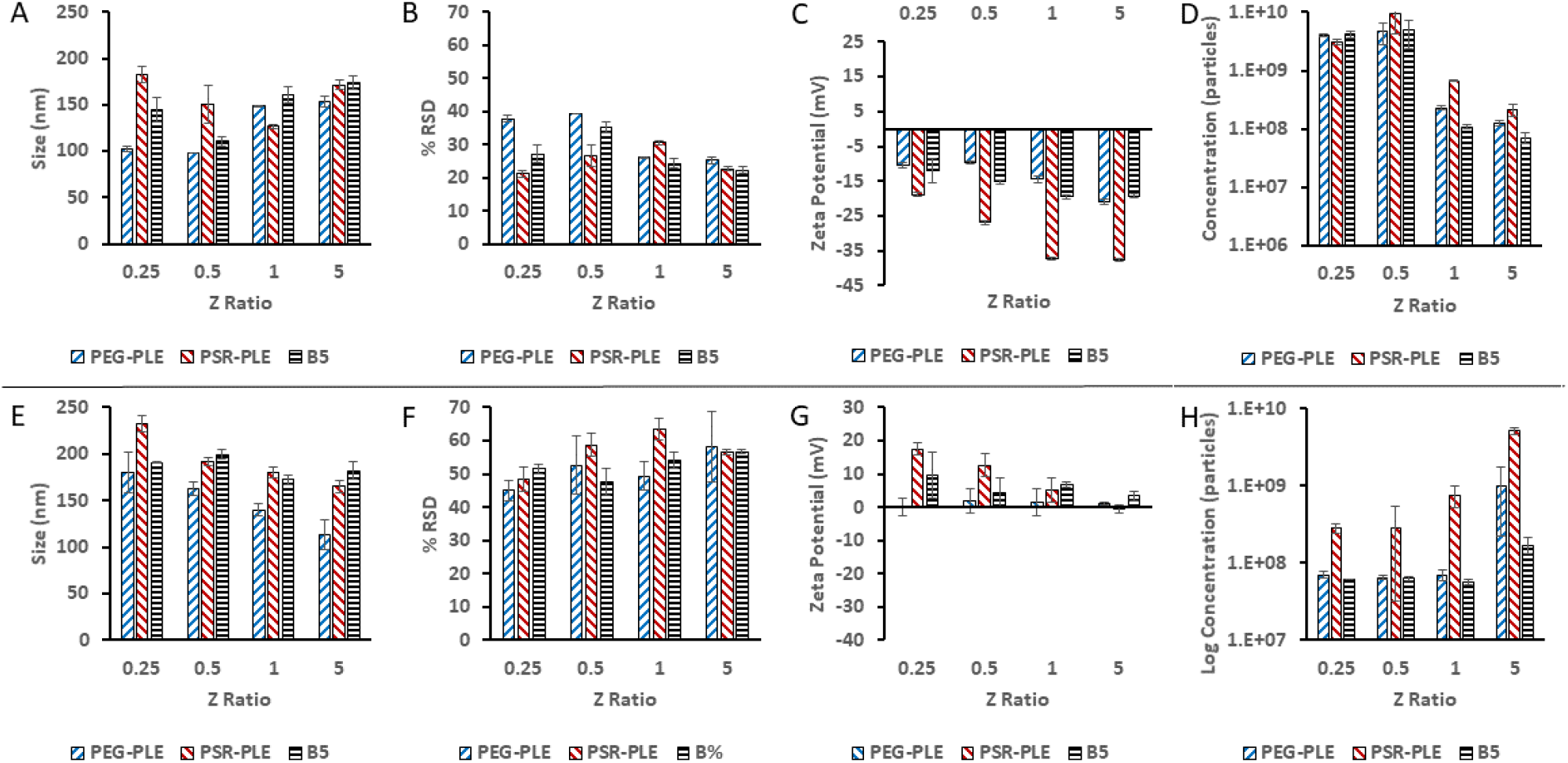
Particle characterization by (A, E) size, (B, F) dispersity, (C, G) zeta potential, and (D-H) concentration. Nano-BDNF was formulated in **(A-D)** 10 mM sodium phosphate, pH 7.4 and **(E-F)** 10 mM sodium acetate, pH 5.0 at room temperature using PEG-PLE, PSR-PLE or B5. Values are mean ± SEM

The self-assembly and overcharging effects in nano-BDNF must be dependent on the degree of ionization of the reacting anionic block copolymer. To test this, we conducted the same experiment at pH 5.0 (**Fig. 8E-H**) using a 10 mM sodium acetate buffer, where the polyanions have lower ionization degree (supplementary **Fig. S16, Table S1**). In this case, the complexes formed at Z_-/+_ = 0.25 appeared to have positive charge, which gradually decreased as the amount of the anionic copolymer increased (**Fig. 8G**). Interestingly, there was an increase in the concentration of the nanoparticles, most distinct in the cases of PEG-PLE and PSR-PLE, which suggests transition to PICs with lower molecular mass and aggregation numbers, i.e., opposite to what is seen at pH 7.4.

### 3.6. Thermodynamic Analysis of BDNF-Polyanion Association

To further investigate the phenomena observed in the previous experiment, we used ITC to measure the heat released as we added copolymer to BDNF. Our experiments were designed to keep the BDNF concentration constant as the anionic block copolymer increased step by step. The copolymer concentrations were adjusted to keep the number of polyacid repeating units uniform across different copolymers. Thus, the x-axis in our thermograms correlates to the Z_-/+_ ratio rather than the molar ratio. For clarity, Z_-/+_ ratio equal 1 corresponds to the polyanion/BDNF molar ratio 0.88 for both PEG-PLE and PSR-PLE and 1.63 for B5 due to the differences in the degrees of polymerization of the polyanion blocks of these copolymers (**Fig. 2**). Addition of each copolymer dropwise to a well containing BDNF yielded exothermic reactions (**Fig. 9**). Notably, none of the integrated ITC curves had a simple sigmoidal shape that would be expected for a simple single set of identical sites (SSIS) model for the molecules binding [50]. In fact, for each of the copolymers there was a sharp minimum observed at Z_-/+_ between 0.25 and 0.5. The heat effects level off to the baseline at Z_-/+_ between 0.5 and 0.8. Notably, the pattern of heat release is like what might be expected of multiple ligands binding a transition metal [50]. In our system, this curve might be due to initial formation of nascent PICs formed at low Z_-/+_ which further self-assemble into mature Nano-BDNF nanoparticle as more block copolymer is added.

**Figure 9.**
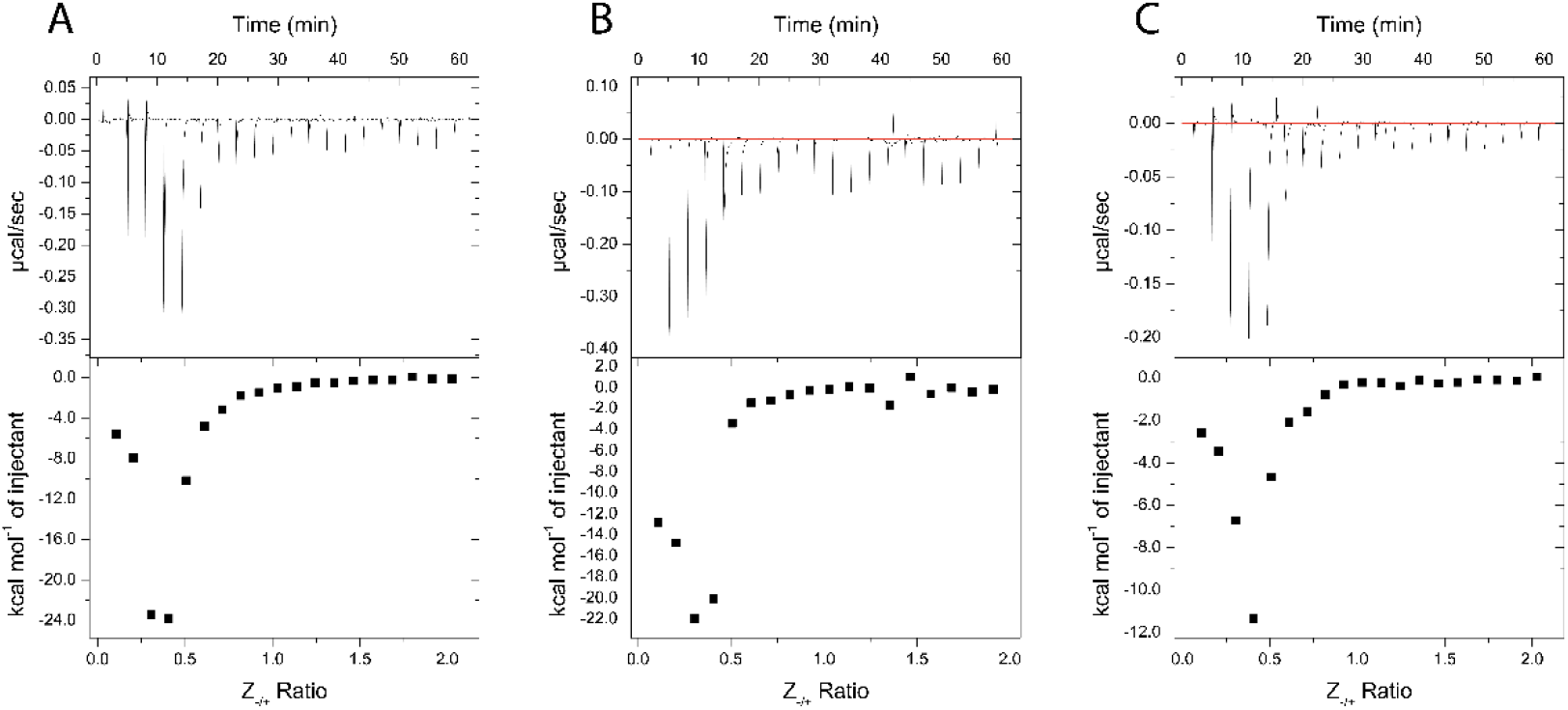
ITC thermograms of Nano-BDNF formation in 10 mM sodium phosphate buffer, pH 7.4. The holding cell was filled with 200 μL of 10 μM BDNF at 25°C. **(A)** 88 μM PEG-PLE **(B)** 88 μM PSR-PLE, and **(C)** 162 μM B5 were injected in 2 μL increments with a 3 min dwell time between each injection. Top: time dependence of the heat supplied to the sample cell for each injection of the block copolymer into the BDNF solution. Bottom - integrated ITC curves for the block copolymer binding with BDNF as a function of the charge ratio Z-/+.

To further validate the relationship between these processes and the charge ratio of the reacting components in the mixture, we again decreased the pH value of the system, keeping all other parameters the same. Although the overall shape of the curves did not change, at pH 5.0 the minimum in integrated ITC curves shifted to higher Z_-/+_ between 0.5 and 0.8 and the saturation region to Z_-/+_ between 1.5 and 2.0 (**Fig. 10**, see also supplementary **Table S2** for comparison of plot features at different pH). The polyanionic blocks (PLE and PPaOx) in our block copolymers undergo ionization in the range from pH 4 to pH 8 (supplementary **Fig. S16, Table S1**). The shift of the position of the minimum in integrated ITC curves is consistent with a decrease in the amount of ionized carboxylic groups in the reacting polyacids that takes place at lower pH, and is indicative of the electrostatic nature of the polyion coupling reactions in this Z_-/+_ range. Interestingly, at pH 5.8 no such shift in the integrated ITC curves minimum and level off regions was observed (supplementary **Figure S18**), which is probably explained by cooperative stabilization of the PICs as will be discussed below. The overall magnitude of the exothermic heat effects seen at the integrated ITC curves minima differed for block copolymers and were notably smaller for B5, for which the heat release dramatically decreased at lower pH (supplementary **Table S2**).

**Figure 10.**
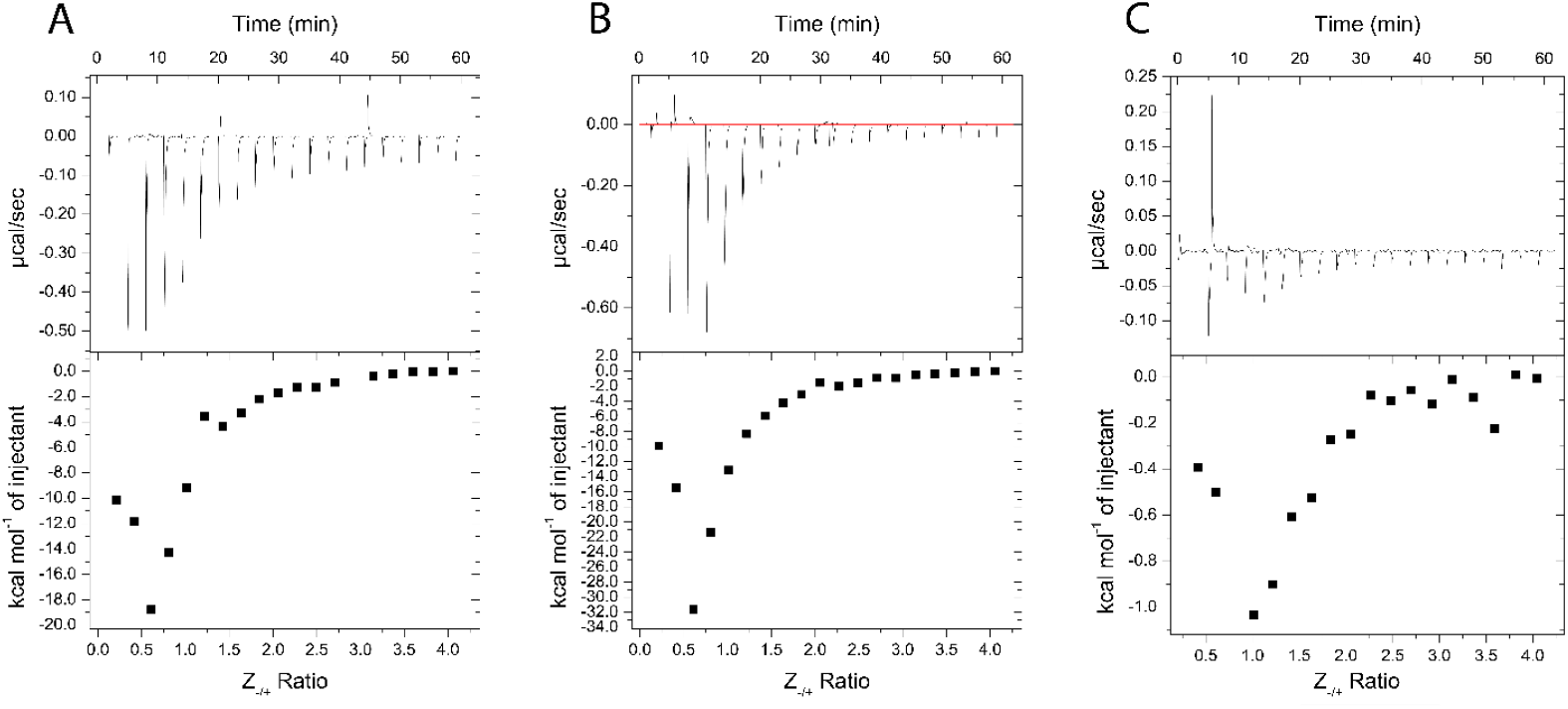
ITC thermograms of Nano-BDNF formation in 10 mM sodium acetate buffer, pH 5.0. The holding cell was filled with 200 μL of 10 μM BDNF in at 25°C. (A) 88 μM PEG-PLE **(B)** 88 μM PSR-PLE, and **(C)** 162 μM B5 were injected in 2 μL increments with a 3 min dwell time between each injection. Top: time dependence of the heat supplied to the sample cell for each injection of the block copolymer into the BDNF solution. Bottom - integrated ITC curves for the block copolymer binding with BDNF as a function of the charge ratio Z-/+.

### 3.7. Stimulation of the TrkB Pathway

The potential therapeutic application of BDNF is premised on the effective stimulation of the TrkB kinase pathway which effects neuropotentiation, neuron survival, and repair [14,43]. Nanoformulation entails the possibility of making BDNF unavailable to the receptor TrkB. We applied Nano-BDNF to NIH 3T3 cells stably transfected with TrkB and measured the phosphorylation of a downstream kinase, extracellular signal-regulated kinase 1/2 (Erk). BDNF, Nano-BDNF B5 Z±=2, and Nano-BDNF PSR-PLE Z_±_=5 stimulated the phosphorylation of Erk compared to the total Erk present in the cell (**Fig 11**). The availability of BDNF was acceptable and indicates that neither of our novel Nano-BDNF formulations inhibits TrkB signaling.

**Figure 11.**
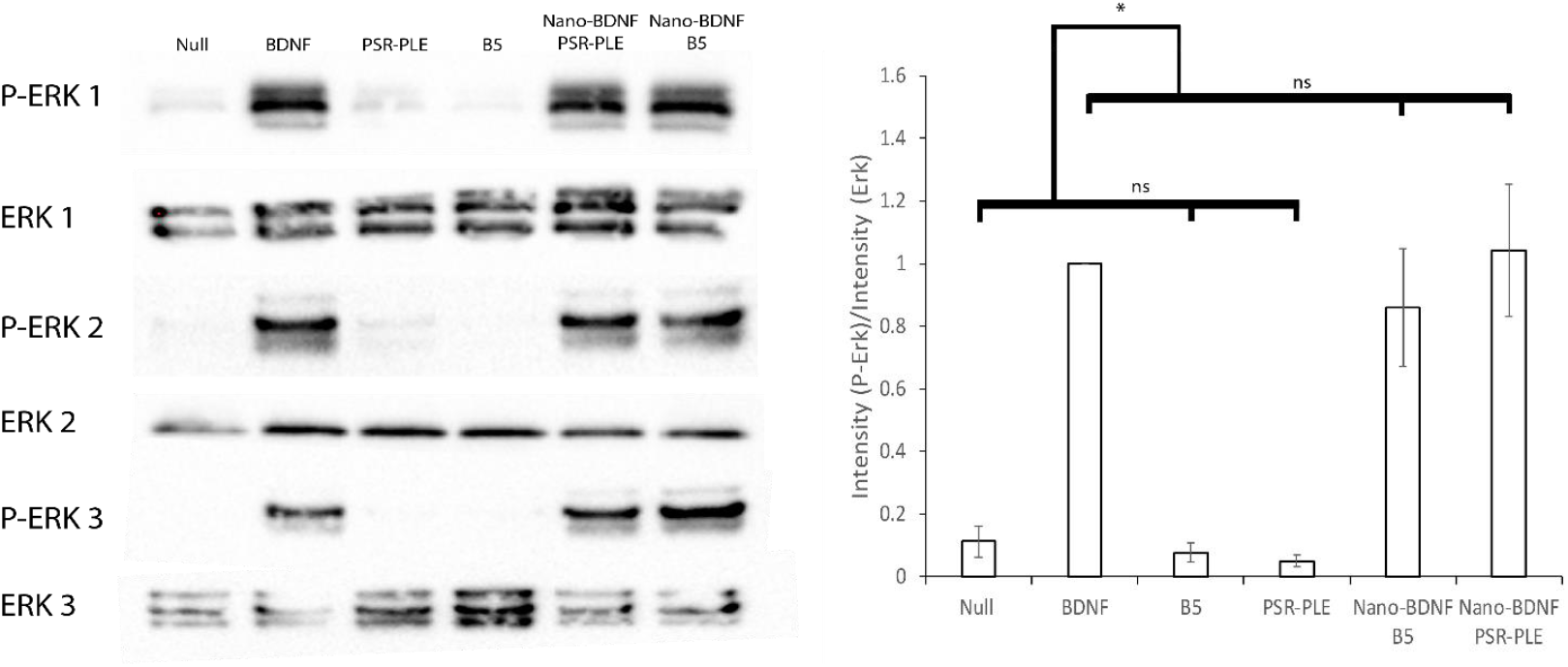
Stimulation of the TrkB Pathway by application of BDNF and Nano-BDNF formulations to NIH 3T3 TrkB cells. For each experiment (1-3), cultured cells were stimulated with 500 ng/mL of BDNF, equivalent amount of nano-BDNF formulated in media at Z_-/+_ = 5, or polymer alone. **(A)** Erk or P-Erk was detected using α-Erk or α-P-Erk via western blot. **(B)** The adjusted total lane volume (measured in signal intensity) of each lane on the blots was determined then the ratio of p-ErK/Erk was calculated. All values were normalized to the BDNF sample and averaged. Experiments were done in triplicate from the point of cell culture. Values are mean ± SEM, * p < 0.05. Statistical significance was established using an unpaired two tailed Student’s t-test.

## 4. Discussion

The well-established immunogenicity of PEG and the high frequency of endogenous α-PEG antibodies indicate a dire need for alternative drug encapsulation technologies with comparable or lower immunogenic profiles [3–7]. A recent study identified anaphylaxis caused by the COVID-19 vaccines which was disproportionally prevalent in women, most likely influenced by higher exposure in household products containing PEG [8,9]. Both poly(sarcosine) and poly(methyl-2-oxazolines) have demonstrated physical, protective, and stealth characteristics similar to PEG [19–21,26,27,31,39,42]. Critically, these alternative polymers do not currently have such extensive applications across a number of industries which limits exposure for most of the population. Both poly(sarcosine) and poly(oxazolines) are relatively simple to manufacture and functionalize or are commercially available. Poly(oxazolines) in particular are a broad class of molecules that allow for hydrophobic, charged, and otherwise reactive moieties to be included in polymer structures [19,30,33,51]. Other work has taken advantage of the variety of poly(oxazoline) functional groups including as the stealth portion of the nanoparticles [30,31,33,36,44,52]. Additionally, LCROP allows excellent control over the length and order of blocks which are easily assessed using ^1^H NMR to determine structure as demonstrated in the results section.

The original Nano-BDNF formulation using PEG-PLE, described in Jiang, et al. and Harris, et al., demonstrated promising effects in disease models [10,11]. Indeed, the simple self-assembly made similar nanoparticles especially persuasive to attempt to reformulate. Our candidate molecules with non-PEG protective blocks were suggested by their similarity to original formulation, especially in the context of the associative block. A winnowing strategy eliminated the polymers which yielded poorly formed or inconsistent particles and we found that optimal particles were formed in the same buffer as Nano-BDNF PEG-PLE.

We implemented microfluidic mixing throughout our study to prepare the complexes. The small-scale batch formulation by manual mixing carries the inherent drawback of batch-by-batch variation due to non-uniform mixing. We overcome this limitation by extensive use of microfluidic mixing which, in addition to improving reproducibility, decreases size and narrows dispersity by mixing the components in a controlled manner. Microfluidic mixing has become an increasingly common technique, especially in formulation of lipid nanoparticles [48]. Furthermore, in our experience, the dispersity of PICs is improved through application of this technology and consistency across batches is also improved [11]. Here, our choice of a reusable, simple, cheap, in house designed mixer indicates that even low-level implementation of these technologies can improve the consistency and quality of formulations.

Nanoformulation carries inherent risk of deactivation of an intended therapy, especially for signaling molecules which must access and bind to a target receptor. Here we demonstrate that despite inclusion of the BDNF into the PICs at the excess of the copolymers, both our reformulated Nano-BDNF variants remain active with respect to TrkB signaling pathway in cells. This has been previously reported by us for Nano-BDNF PEG-PLE and the mechanism was demonstrated [11]. Specifically, we have previously shown that while the complexes remain stable in the presence of salt and serum proteins, the TrkB receptor can readily abstract the BDNF molecule, presumably due to higher affinity to the BDNF active site compared to the anionic block copolymer. In this case, TrkB can access and bind BDNF which yields a signaling cascade that we monitor via Erk phosphorylation. Moreover, based on prior molecular modeling analysis [11], the polyanion binds to the same active site as the TrkB thereby competitively masking this site in the absence of the specific receptor. Therefore, this class of PICs is particularly well suited for delivery of BDNF and possibly other neutrophins. Non-covalent interactions allow some reorganization and may enable a receptor with strong, specific binding to access the signaling molecule. Conversely, the preferential binding with the anionic blocks of the copolymers protects from opsonization [10,11].

Our study also provides new insight on the complexation of the BDNF with anionic block copolymers. Using the highly efficient, multi-utility nanoparticle tracking technique, we confirm the formation of the PICs at the deficiency of the anionic block copolymers. Based on the patterns of the particle zeta potential and concentration, we propose that these complexes form initially as the nascent species that reorganize into mature Nano-BDNF particles as the amounts of the added anionic block copolymers (Z_-/+_ ratio) increase. An independent ITC experiment confirms the distinct processes at lower and higher Z_-/+_ ratios. The processes of polymer coupling at the block copolymer deficiency at lower Z_-/+_ are strongly exothermic which results in appearance of the deep minima in the integrated curve. This exothermic effect vanished as the amount of the block copolymer increased at higher Z_-/+_. The data from our ITC experiments were not easily fit by SSIS models or a two-step model like DNA condensation [50,53,54]. However, the ITC pattern observed for complexation of net cationic BDNF with our polyanions appears to qualitatively mirror that previously observed for the binding of DNA with polycations as reported by us and Kataoka [53,55].

There are, however, notable differences. First, our systems display strongly exothermic effects while the DNA and polycation binding is endothermic. Second, the magnitude of our effect, although varying for different polyanions, is at least one order of magnitude greater than the effects observed for DNA and polycation binding. The small enthalpic effects observed for DNA and polycation binding are typical for entropy driven processes due to the counterion release [55], and could also be compounded by the conformational transitions due to the DNA condensation [53]. In our case, BDNF displays 44 unique, positively charged Lys and Arg residues on its exterior and based on the prior molecular modeling binding of the polycarboxylic acid to it involves formation of multiple hydrogen bonds [11]. The hydrogen bond formation is an exothermic process which is consistent with our observation [56].

Unsurprisingly, the complexation is dependent on pH which alters ionization of the polyacid. In the supplement we provided the titration results and ionization characteristics of all three anionic block copolymer used in our work. As we decrease the pH to 5.0, the degree of ionization decreases dramatically, and the characteristics of the complexes change. By the nanoparticle tracking technique we observe formation of positively charged species at low amounts of the added anionic block copolymers (Z_-/+_ ratio). We also observe changes in the ITC patterns where the amount of the block copolymer required in the exothermic binding increases. The greatest effects appear with the poly(oxazoline) block copolymer B5, which has the highest value of pK_a_. This suggests strong effect of the polyanion ionization in the formation of Nano-BDNF complexes. However, we would like to point out that the direct quantitative relation of the degree of ionization of the weak polyelectrolytes to polyion complexation should be done with caution because of the cooperative stabilization of the ionized forms [57]. Specifically, in the discussed pH range the BDNF molecule is positively charged, and binding to it can stabilize the ionized forms of the polycarboxylic acids even at low pH (a phenomenon we cannot demonstrate here by potentiometric titration because of prohibitive costs). This may explain why we do not see much change in the ITC results at pH 5.8. Most importantly, our current findings suggest that by carefully selecting pH during the manufacturing processes, the properties of polyelectolyte formulations of neurotrophins can be optimized. Overall, deeper understanding of the physics which dictate nanoparticle formation and how varying conditions affect formulation can yield valuable insight to how future polyelectrolyte complex formulations are designed or reformulated.

## 5. Conclusions

We demonstrate the suitability of two novel polymers to substitute for PEG-PLE in reformulations of Nano-BDNF that are free of PEG. Particles of similar size and properties are formed through simple, microfluidic mixing. Overall, we demonstrate that by preserving the specific attributes of the associative and protective blocks, we can produce nanoparticles that behave in a similar manner. The new results are instrumental for the design of future formulations of neurotrophins for clinical use.

## Supporting information

supplement for article

## Author Contributions

Conceptualization, James Fay, Chaemin Lim, Elena Batrakova and Alexander Kabanov; Data curation, James Fay and Chaemin Lim; Formal analysis, James Fay, Chaemin Lim and Alexander Kabanov; Funding acquisition, Elena Batrakova and Alexander Kabanov; Investigation, James Fay and Anna Finkelstein; Methodology, James Fay, Chaemin Lim, Anna Finkelstein, Elena Batrakova and Alexander Kabanov; Project administration, Alexander Kabanov; Resources, Elena Batrakova and Alexander Kabanov; Supervision, Elena Batrakova and Alexander Kabanov; Validation, James Fay, Chaemin Lim and Alexander Kabanov; Visualization, James Fay; Writing – original draft, James Fay and Alexander Kabanov; Writing – review & editing, James Fay, Elena Batrakova and Alexander Kabanov.

## Acknowledgments

We would like to thank Dr. David Kaplan (Department of Molecular Genetics, University of Toronto, Canada) for his kind gift of NIH 3T3 TrkB cells which continues to bear fruit, Dr. Ashutosh Tripathy for his aid and instruction in ITC experiments and analysis, Dr. Elizabeth Wayne, Ms. Yuling Zhao, and Mr. Matthew Haney for their aid and instruction in cell culture, Dr. Dina Yamalaeva and Dr. Natasha Vinod for production of the microfluidic mixers, Ms. Jubina Bregu for her administrative support and advice, Dr. Yuhang Jiang for his kind encouragement and base work for the project, and Ms. Bethany Boring Esq. for proofreading the final draft.

## Data Availability

Data are available from authors upon reasonable request.

## References

1. Gref, R.; Minamitake, Y.; Peracchia, M.T.; Torchilin, V.; Langer, R.; Peracchia, T.; Trubetskoy, V.; Torchilin, V.; Langerllf, R. Biodegradable Long-Circulating Polymeric Nanospheres. Science 1994, 263, 1600–1603.

2. Armstrong, J.K. PEGylated Protein Drugs: Basic Science and Clinical Applications; Birkhauser: Basel, Switzerland, 2009; ISBN 9783764386788.

3. Ishida, T.; Masuda, K.; Ichikawa, T.; Ichihara, M.; Irimura, K.; Kiwada, H. Accelerated Clearance of a Second Injection of PEGylated Liposomes in Mice. Int. J. Pharm. 2003, 255, 167–174, doi:10.1016/S0378-5173(03)00085-1.

4. Yang, Q.; Jacobs, T.M.; McCallen, J.D.; Moore, D.T.; Huckaby, J.T.; Edelstein, J.N.; Lai, S.K. Analysis of Pre-Existing IgG and IgM Antibodies against Polyethylene Glycol (PEG) in the General Population. Anal. Chem. 2016, 88, 11804–11812, doi:10.1021/acs.analchem.6b03437.

5. Risma, K.A.; Edwards, K.M.; Hummell, D.S.; Little, F.F. Potential Mechanisms of Anaphylaxis to COVID-19 MRNA Vaccines. J. Allergy Clin. Immunol. 2019, 147, 2075–2082.e2, doi:10.1016/j.jaci.2021.04.002.

6. Haddad, H.F.; Burke, J.A.; Scott, E.A.; Ameer, G.A. Clinical Relevance of Pre-Existing and Treatment-Induced Anti-Poly ( Ethylene Glycol ) Antibodies. Regen. Eng. Transl. Med. 2021, 8, 32–42, doi:10.1007/s40883-021-00198-y.

7. Moghimi, S.M. Allergic Reactions and Anaphylaxis to LNP-Based COVID-19 Vaccines. Mol. Ther. 2021, 29, 898–900, doi:10.1016/j.ymthe.2021.01.030.

8. Warren, C.M.; Snow, T.T.; Lee, A.S.; Shah, M.M.; Heider, A.; Blomkalns, A.; Betts, B.; Buzzanco, A.S.; Gonzalez, J.; Chinthrajah, R.S.; et al. Assessment of Allergic and Anaphylactic Reactions to MRNA COVID-19 Vaccines with Confirmatory Testing in a US Regional Health System. JAMA Netw. Open 2021, 4, 1–15, doi:10.1001/jamanetworkopen.2021.25524.

9. Somiya, M.; Mine, S.; Yasukawa, K.; Ikeda, S. Sex Differences in the Incidence of Anaphylaxis to LNP-MRNA COVID-19 Vaccines. Vaccine 2021, 39, 3313–3314.

10. Harris, N.M.; Ritzel, R.; Mancini, N.; Jiang, Y.; Yi, X.; Manickam, D.S.; Banks, W.A.; Kabanov, A. V.; McCullough, L.D.; Verma, R. Nano-Particle Delivery of Brain Derived Neurotrophic Factor after Focal Cerebral Ischemia Reduces Tissue Injury and Enhances Behavioral Recovery. Pharmacol. Biochem. Behav. 2016, 150–151, 48–56, doi:10.1016/j.pbb.2016.09.003.

11. Jiang, Y.; Fay, J.M.; Poon, C.D.; Vinod, N.; Zhao, Y.; Bullock, K.; Qin, S.; Manickam, D.S.; Yi, X.; Banks, W.A.; et al. Nanoformulation of Brain-Derived Neurotrophic Factor with Target Receptor-Triggered-Release in the Central Nervous System. Adv. Funct. Mater. 2018, 28, 1–11, doi:10.1002/adfm.201703982.

12. Nagahara, A.H.; Merrill, D.A.; Coppola, G.; Tsukada, S.; Schroeder, B.E.; Shaked, G.M.; Wang, L.; Blesch, A.; Kim, A.; Conner, J.M.; et al. Neuroprotective Effects of Brain-Derived Neurotrophic Factor in Rodent and Primate Models of Alzheimer’s Disease. Nat. Med. 2009, 15, 331–337, doi:10.1038/nm.1912.

13. Tsukahara, T.M.D.; Takeda, M.M..; Shimohama, S.M..; Ohara, O.P..; Hashimoto, N.M.. Effects of Brain-Derived Neurotrophic Factor on 1-Methyl-4-Phenyl-1,2,3,6-Tetrahydropyridine-Induced Parkinsonism in Monkeys. Neurosurgery 1995, 37, 733–741, doi:10.1227/00006123-199510000-00018.

14. Zhao, H.; Alam, A.; San, C.-Y.; Eguchi, S.; Chen, Q.; Lian, Q.; Ma, D. Molecular Mechanisms of Brain-Derived Neurotrophic Factor in Neuro-Protection: Recent Developments. Brain Res. 2017, 1665, 1–21, doi:10.1016/j.brainres.2017.03.029.

15. Pan, W.; Banks, W.A.; Fasold, M.B.; Bluth, J.; Kastin, A.J. Transport of Brain-Derived Neurotrophic Factor across the Blood-Brain Barrier. Neuropharmacology 1998, 37, 1553–1561, doi:10.1016/S0028-3908(98)00141-5.

16. Veronese, F.M.; Mero, A.; Pasut, G. Protein PEGylation, Basic Science and Biological Applications. In PEGylated Protein Drugs: Basic Science and Clinical Applications; Veronese, F.M., Ed.; Birkhäuser Basel: Basel, 2009; pp. 11–31 ISBN 978-3-7643-8679-5.

17. Armstrong, J.K. The Occurrence, Induction, Specificity and Potential Effect of Antibodies against Poly(Ethylene Glycol). In PEGylated Protein Drugs: Basic Science and Clinical Applications; Veronese, F.M., Ed.; Birkhäuser Basel: Basel, 2009; pp. 147–168 ISBN 978-3-7643-8679-5.

18. Abuchowski, A.; Es, T. Van; Palczuk, N.C.; Davis, F.F. Alteration of Immunological Properties of Bovine Serum Albumin by Covalent Attachment of Polyethylene Glycol *. J. Biol. Chem. 1977, 252, 3578–3581, doi:10.1016/S0021-9258(17)40291-2.

19. Konradi, R.; Acikgoz, C.; Textor, M. Polyoxazolines for Nonfouling Surface Coatings - A Direct Comparison to the Gold Standard PEG. Macromol. Rapid Commun. 2012, 33, 1663–1676, doi:10.1002/marc.201200422.

20. Son, K.; Ueda, M.; Taguchi, K.; Maruyama, T.; Takeoka, S. Evasion of the Accelerated Blood Clearance Phenomenon by Polysarcosine Coating of Liposomes. J. Control. Release 2020, 322, 209–216, doi:10.1016/j.jconrel.2020.03.022.

21. Luxenhofer, R.; Han, Y.; Schulz, A.; Tong, J.; He, Z.; Kabanov, A. V; Jordan, R. Poly(2-Oxazoline)s as Polymer Therapeutics. Macromol. Rapid Commun. 2012, 33, 1613–1631.

22. Li, Y.; Bronich, T.K.; Chelushkin, P.S.; Kabanov, A. V. Dynamic Properties of Block Ionomer Complexes with Polyion Complex Cores. Macromolecules 2008, 41, 5863–5868, doi:10.1021/ma702671w.

23. He, Z.; Miao, L.; Jordan, R.; S-Manickam, D.; Luxenhofer, R.; Kabanov, A. V. A Low Protein Binding Cationic Poly(2-Oxazoline) as Non-Viral Vector. Macromol. Biosci. 2015, 15, 1004–1020, doi:10.1002/mabi.201500021.

24. Tong, J.; Yi, X.; Luxenhofer, R.; Banks, W.A.; Jordan, R.; Zimmerman, M.C.; Kabanov, A. V Conjugates of Superoxide Dismutase 1 with Amphiphilic Poly(2-Oxazoline) Block Copolymers for Enhanced Brain Delivery: Synthesis, Characterization and Evaluation in Vitro and in Vivo. Mol. Pharm. 2013, 10, 360–377, doi:10.1021/mp300496x.

25. Jiang, Y.; Brynskikh, A.M.; S-Manickam, D.; Kabanov, A. V. SOD1 Nanozyme Salvages Ischemic Brain by Locally Protecting Cerebral Vasculature. J. Control. Release 2015, 213, 36–44, doi:10.1016/j.jconrel.2015.06.021.

26. Barz, M.; Luxenhofer, R.; Zentel, R.; Vicent, M.J. Overcoming the PEG-Addiction: Well-Defined Alternatives to PEG, from Structure–Property Relationships to Better Defined Therapeutics. Polym. Chem. 2011, 2, 1900–1918, doi:10.1039/C0PY00406E.

27. Gao, S.; Holkar, A.; Srivastava, S. Protein – Polyelectrolyte Complexes and Micellar Assemblies. Polymers (Basel). 2019, 11, doi:10.3390/polym11071097.

28. Miyata, K.; Nishiyama, N.; Kataoka, K. Rational Design of Smart Supramolecular Assemblies for Gene Delivery: Chemical Challenges in the Creation of Artificial Viruses. Chem. Soc. Rev. 2012, 41, 2562–2574, doi:10.1039/C1CS15258K.

29. He, Z.; Wan, X.; Schulz, A.; Bludau, H.; Dobrovolskaia, M.A.; Stern, S.T.; Montgomery, S.A.; Yuan, H.; Li, Z.; Alakhova, D.; et al. Biomaterials A High Capacity Polymeric Micelle of Paclitaxel: Implication of High Dose Drug Therapy to Safety and in Vivo Anti-Cancer Activity. Biomaterials 2016, 101, 296–309, doi:10.1016/j.biomaterials.2016.06.002.

30. Vinod, N.; Hwang, D.; Azam, S.H.; Van Swearingen, A.E.D.; Wayne, E.; Fussell, S.C.; Sokolsky-Papkov, M.; Pecot, C. V; Kabanov, A. V High-Capacity Poly(2-Oxazoline) Formulation of TLR 7/8 Agonist Extends Survival in a Chemo-Insensitive, Metastatic Model of Lung Adenocarcinoma. Sci. Adv. 2020, 6, doi:10.1126/sciadv.aba5542.

31. Seo, Y.; Schulz, A.; Han, Y.; He, Z.; Bludau, H.; Wan, X.; Tong, J.; Bronich, T.K.; Sokolsky, M.; Luxenhofer, R.; et al. Poly(2-Oxazoline) Block Copolymer Based Formulations of Taxanes: Effect of Copolymer and Drug Structure, Concentration, and Environmental Factors. Polym. Adv. Technol. 2015, 26, 837–850, doi:10.1002/pat.3556.

32. He, Z.; Schulz, A.; Wan, X.; Seitz, J.; Bludau, H.; Alakhova, D.Y.; Darr, D.B.; Perou, C.M.; Jordan, R.; Ojima, I.; et al. Poly ( 2-Oxazoline ) Based Micelles with High Capacity for 3rd Generation Taxoids: Preparation, in Vitro and in Vivo Evaluation. J. Control. Release 2015, 208, 67–75, doi:10.1016/j.jconrel.2015.02.024.

33. Hwang, D.; Ramsey, J.D.; Makita, N.; Sachse, C.; Jordan, R.; Sokolsky-papkov, M.; Kabanov, A. V Novel Poly ( 2-Oxazoline ) Block Copolymer with Aromatic Heterocyclic Side Chains as a Drug Delivery Platform. J. Control. Release 2019, 307, 261–271, doi:10.1016/j.jconrel.2019.06.037.

34. Hwang, D.; Dismuke, T.; Tikunov, A.; Rosen, E.P.; Kagel, J.R.; Ramsey, J.D.; Lim, C.; Zamboni, W.; Kabanov, A. V; Gershon, T.R.; et al. Poly ( 2-Oxazoline ) Nanoparticle Delivery Enhances the Therapeutic Potential of Vismodegib for Medulloblastoma by Improving CNS Pharmacokinetics and Reducing Systemic Toxicity. Nanomedicine Nanotechnology, Biol. Med. 2021, 32, 102345, doi:10.1016/j.nano.2020.102345.

35. Wan, X.; Beaudoin, J.J.; Vinod, N.; Min, Y.; Makita, N.; Bludau, H.; Jordan, R.; Wang, A.; Sokolsky, M.; Kabanov, A. V Biomaterials Co-Delivery of Paclitaxel and Cisplatin in Poly ( 2-Oxazoline ) Polymeric Micelles: Implications for Drug Loading, Release, Pharmacokinetics and Outcome of Ovarian and Breast Cancer Treatments. Biomaterials 2019, 192, 1–14, doi:10.1016/j.biomaterials.2018.10.032.

36. Hwang, D.; Ramsey, J.D.; Kabanov, A. V Polymeric Micelles for the Delivery of Poorly Soluble Drugs: From Nanoformulation to Clinical Approval. Adv. Drug Deliv. Rev. 2020, 156, 80–118, doi:10.1016/j.addr.2020.09.009.

37. Tong, J.; Yi, X.; Luxenhofer, R.; Banks, W.A.; Jordan, R.; Zimmerman, M.C.; Kabanov, A. V. Conjugates of Superoxide Dismutase 1 with Amphiphilic Poly(2-Oxazoline) Block Copolymers for Enhanced Brain Delivery: Synthesis, Characterization and Evaluation in Vitro and in Vivo. Mol. Pharm. 2013, 10, 360–377, doi:10.1021/mp300496x.

38. Lau, K.H.A. Peptoids for Biomaterials Science. Biomater. Sci. 2014, 2, 627, doi:10.1039/c3bm60269a.

39. Cheung, D.L.; Hang, K.; Lau, A. Atomistic Study of Zwitterionic Peptoid Antifouling Brushes. 2019, doi:10.1021/acs.langmuir.8b01939.

40. Huesmann, D.; Sevenich, A.; Weber, B.; Barz, M. A Head-to-Head Comparison of Poly(Sarcosine) and Poly(Ethylene Glycol) in Peptidic, Amphiphilic Block Copolymers. Polym. (United Kingdom) 2015, 67, 240–248, doi:10.1016/j.polymer.2015.04.070.

41. Chan, B.A.; Xuan, S.; Li, A.; Simpson, J.M.; Sternhagen, G.L.; Yu, T.; Darvish, O.A. Polypeptoid Polymers: Synthesis, Characterization, and Properties. 2018, 1–25, doi:10.1002/bip.23070.

42. Hu, Y.; Hou, Y.; Wang, H.; Lu, H. Polysarcosine as an Alternative to PEG for Therapeutic Protein Conjugation. Bioconjug. Chem. 2018, 29, 2232–2238, doi:10.1021/acs.bioconjchem.8b00237.

43. Soppet, D.; Escandon, E.; Maragos, J.; Middlemas, D.S.; Raid, S.W.; Blair, J.; Burton, L.E.; Stanton, B.R.; Kaplan, D.R.; Hunter, T. The Neurotrophic Factors Brain-Derived Neurotrophic Factor and Neurotrophin-3 Are Ligands for the TrkB Tyrosine Kinase Receptor. Cell 1991, 65, 895–903, doi:10.1093/glycob/cwn084.

44. Luxenhofer, R.; Schulz, A.; Roques, C.; Li, S.; Bronich, T.K.; Batrakova, E. V; Jordan, R.; Kabanov, A. V Doubly-Amphiphilic Poly(2-Oxazoline)s as High Capacity Delivery Systems for Hydrophobic Drugs. Biomaterials 2011, 31, 4972–4979, doi:10.1016/j.biomaterials.2010.02.057.Doubly-Amphiphilic.

45. Kabanov, V.A. Polyelectrolyte Complexes in Solution and in Bulk. Russ. Chem. Rev. 2005, 74, 3–20, doi:10.1070/RC2005v074n01ABEH001165.

46. Chelushkin, P.S.; Lysenko, E.A.; Bronich, T.K.; Eisenberg, A.; Kabanov, V.A.; Kabanov, A. V. Polyion Complex Nanomaterials from Block Polyelectrolyte Micelles and Linear Polyelectrolytes of Opposite Charge. 2. Dynamic Properties. J. Phys. Chem. B 2008, 112, 7732–7738, doi:10.1021/jp8012877.

47. Lu, M.; Ozcelik, A.; Grigsby, C.L.; Zhao, Y.; Guo, F.; Leong, K.W.; Huang, T.J. Microfluidic Hydrodynamic Focusing for Synthesis of Nanomaterials. Nano Today 2016, 11, doi:http://dx.doi.org/10.1016/j.nantod.2016.10.006.

48. Guimaraes Sa Correia, M.; Briuglia, M.L.; Niosi, F.; Lamprou, D.A. Microfluidic Manufacturing of Phospholipid Nanoparticles: Stability, Encapsulation Efficacy, and Drug Release. Int. J. Pharm. 2017, 516, 91–99, doi:10.1016/j.ijpharm.2016.11.025.

49. Burns, M.L.; Malott, T.M.; Metcalf, K.J.; Hackel, B.J.; Chan, J.R.; Shusta, E. V. Directed Evolution of Brain-Derived Neurotrophic Factor for Improved Folding and Expression in Saccharomyces Cerevisiae. Appl. Environ. Microbiol. 2014, 80, 5732–5742, doi:10.1128/AEM.01466-14.

50. ITC Data Analysis in Origin; 7th ed.; MicroCal LLC: Northhampton, MA, 2004;

51. Ulbricht, J.; Jordan, R.; Luxenhofer, R. On the Biodegradability of Polyethylene Glycol, Polypeptoids and Poly(2-Oxazoline)S. Biomaterials 2014, 35, 4848–4861, doi:10.1016/j.biomaterials.2014.02.029.

52. Koshkina, O.; Westmeier, D.; Lang, T.; Bantz, C.; Hahlbrock, A.; Würth, C.; Resch-Genger, U.; Braun, U.; Thiermann, R.; Weise, C.; et al. Tuning the Surface of Nanoparticles: Impact of Poly(2-Ethyl-2-Oxazoline) on Protein Adsorption in Serum and Cellular Uptake. Macromol. Biosci. 2016, 1287–1300, doi:10.1002/mabi.201600074.

53. Kim, W.; Yamasaki, Y.; Jang, W.D.; Kataoka, K. Thermodynamics of DNA Condensation Induced by Poly(Ethylene Glycol)- Block -Polylysine through Polyion Complex Micelle Formation. Biomacromolecules 2010, 11, 1180–1186, doi:10.1021/bm901305p.

54. Kim, W.; Yamasaki, Y.; Kataoka, K. Development of a Fitting Model Suitable for the Isothermal Titration Calorimetric Curve of DNA with Cationic Ligands. J. Phys. Chem. B 2006, 110, 10919–10925, doi:10.1021/jp057554e.

55. Bronich, T.; Kabanov, A. V.; Marky, L.A. A Thermodynamic Characterization of the Interaction of a Cationic Copolymer with DNA. J. Phys. Chem. B 2001, 105, 6042–6050, doi:10.1021/jp004395k.

56. Friedman, N. Hydrogen Bonding and Heat of Solution. J. Chem. Educ. 1977, 54, 248, doi:10.1021/ed054p248.

57. Kabanov, A. V; Bronich, T.K.; Kabanov, V.A.; Yu, K.; Eisenberg, A. Soluble Stoichiometric Complexes from Poly ( N-Ethyl-4-Vinylpyridinium ) Cations and Poly ( Ethylene Oxide ) -Block-Polymethacrylate Anions. Society 1996, 9297, 6797–6802.

