## supplement for article for "PEG-Free Polyion Complex Nanocarriers for Brain Derived Neurotrophic Factor"

*Article Supplement*

**Supplementary Figures and Results**

*S1.1. Microfluidic mixer*

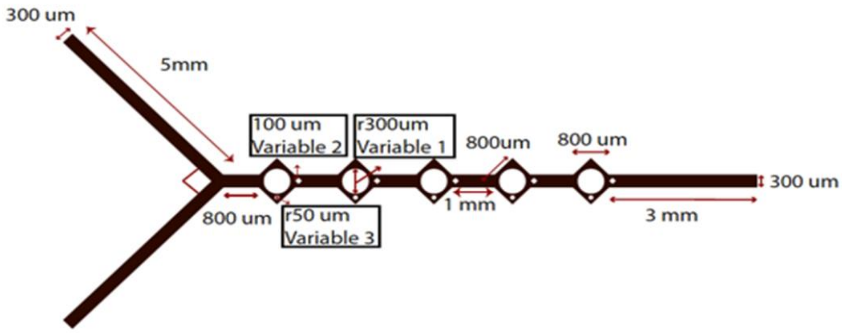


**Figure S1. Schematics showing configuration and dimensions of the mixer used for microfluidics formulation. R**eproduced with permission from Jiang et al.[1]: Much of our work is dependent on the application of an in house designed microfluidic mixer by which we can exert control over the rate of mixing through the device. We introduced this device in the previous report of the precedent work[1] where we observed much improved formulation of Nano-BDNF PEG-PLE.

*S1.2 NMR characterization of B1-B5 polymers*

Here we have the spectra from the $^1$H NMR produced using MestReNova (11.0). These data were used to calculate the polymer block ratios and lengths. Despite minor deviation from expected block lengths, we observe the intended motifs of B1: short-short-short, B2: short-long-short, B3: long-short-long, and B5: long-long-long. It is clear from simple visual inspection that varying the amount of reagents in the reaction allows control over the overall lengths. More precise methods might allow finer control, but we expect a 30 subunit block acts much the same as one of 27 subunits. **Fig. 1** in the main text details peak assignment.

As a brief aside, our B1-5 naming convention may cause some confusion as we omit B4. Originally, these were referenced as batch 1-5 and batch 4 was a failed synthesis. However, in our notes, presentations, etc. we continued to call them by B1, B2…. While the omission of B4 might cause minor confusion, in interest of maintaining data cohesion from notebooks to manuscript, we maintained the prior naming scheme. Later batches were named polymer B5 synthesis 2 rather than batch 7 to avoid confusion.

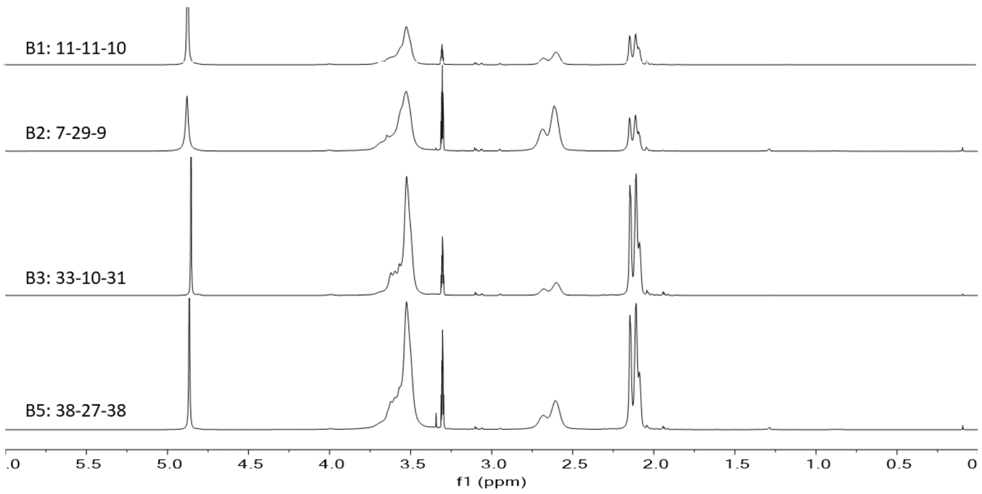


**Figure S2. $^1$H NMR spectra of the four PMeOx-PPaOx-PMeOx polymers**. Polymers were dissolved in deuterated methanol and analyzed in a INOVA 400 NMR machine.

*S1.3 Particle size characterization*

We performed a series of experiments where we measured the size and dispersity of our nanoparticles using the Zetasizer DLS machine. Generally, intensity weighted size is reported for DLS data though the Zetasizer yields number weighted as well. Comparing the number weighted size with Zetasizer or Nanosight data can yield insight whether a few large particles dominate, or the solution consists mostly of free polymer/protein. In no cases did we decide exclusively from the number mean average size, but we did consider it, especially in eliminating B2 during the winnowing process.

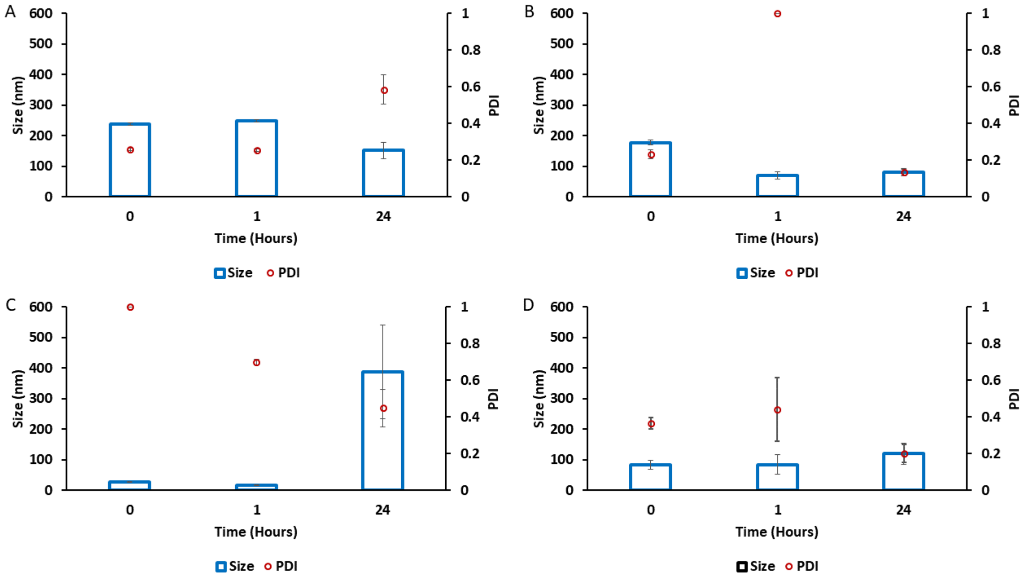


**Figure S3. Intensity weighted particle size and dispersity of Nano-BDNF PSR-PLE.** The complexes were prepared by manual mixing of the solutions of BDNF and PSR-PLE in 10 mM HEPES, pH 7.4 at the $Z_{-/+}$ ratios of (A) 1, (B) 2, (C) 5, and (D) 10. The DLS measurements were carried out for up to 48 hours. Values are mean ± SEM.

.

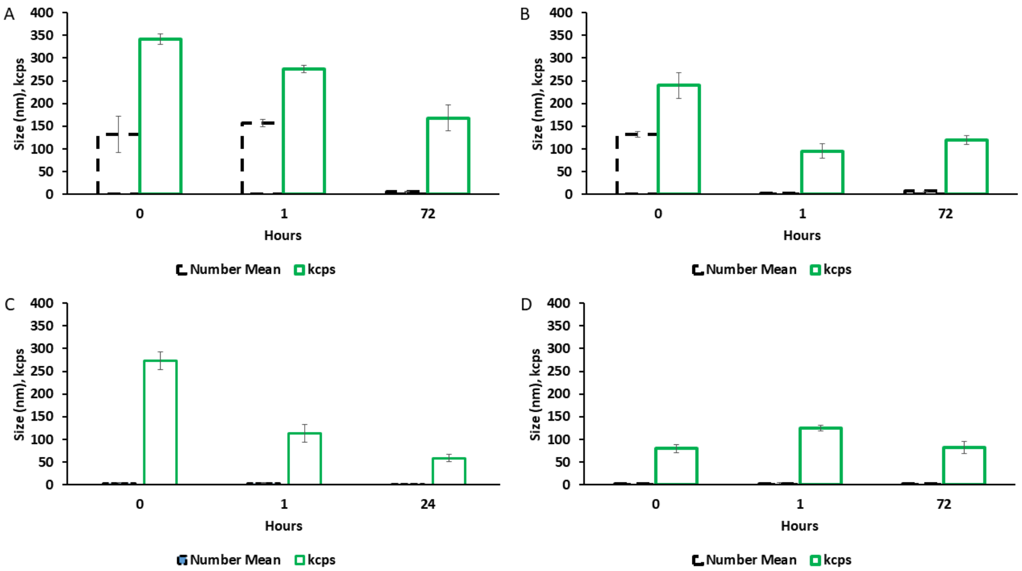


**Figure S4. Number weighted particle size and kilocount per second of Nano-BDNF PSR-PLE measurements in DLS of the same samples as shown Fig. S3.**

*Low ionic strength, minimal buffer for PSR-PLE (Figs. S3 and S4).*

We observed that formation of Nano-BDNF PSR-PLE in 10 mM HEPES, pH 7.4 was inconsistent. While nanoparticles formed at $Z_{-/+}$=1 were acceptable at a short time, the high $Z_{-/+}$ nanoparticles were very variable in size. During this trial we decided to continue development with PSR-PLE due to the similarity of structure with PEG-PLE.

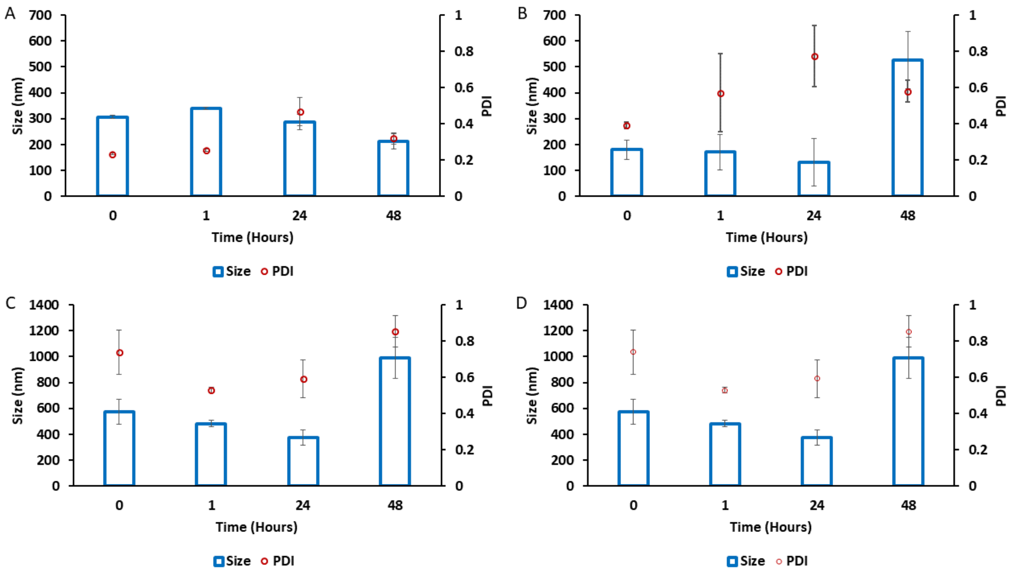


**Figure S5. Intensity weighted particle size and dispersity of Nano-BDNF B1.** The complexes were prepared by manual mixing of the solutions of BDNF and PSR-PLE in 10 mM HEPES, pH 7.4 at the $Z_{-/+}$ ratios of (A) 1, (B) 2, (C) 5, and (D) 10. The DLS measurements were carried out for up to 48 hours. Values are mean ± SEM.

.

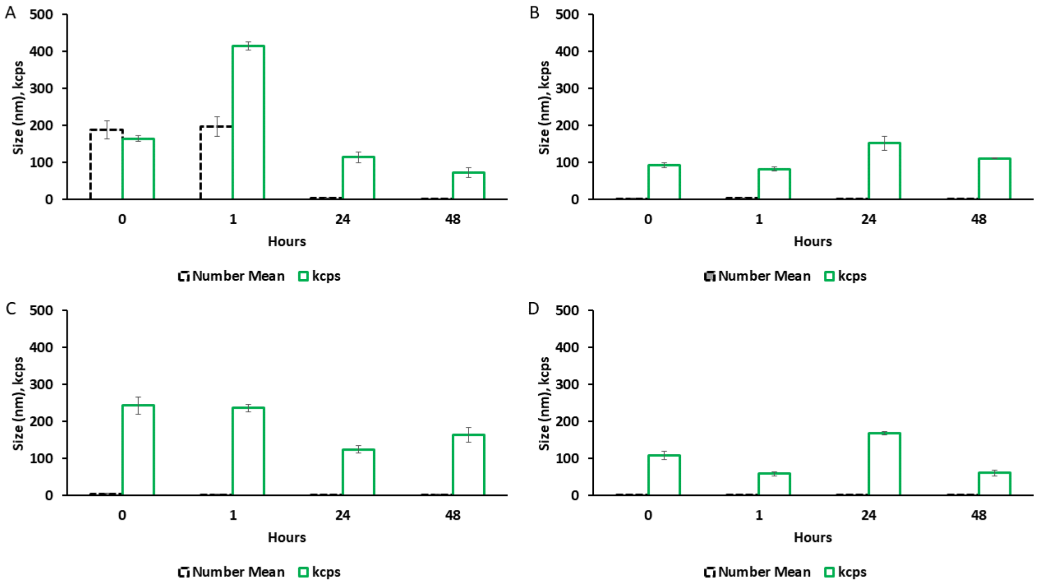


**Figure S6. Number weighted particle size and kilocount per second of Nano-BDNF B1 measurements in DLS of the same samples as shown Fig. S5.**

*Low ionic strength, minimal buffer for B1 (Figs. S5 and S6).*

Nano-BDNF B1 ($PMeOx_{11}$-$PPaOx_{11}$-$PMeOx_{10}$) formulated in 10 mM HEPES, pH 7.4 generally formed large nanoparticles or aggregates. At $Z_{-/+}$ =1 we observed the best results where the size was under 400 nm, but the other $Z_{-/+}$ ratios convinced us to pursue other polymers due to large sizes and high dispersity.

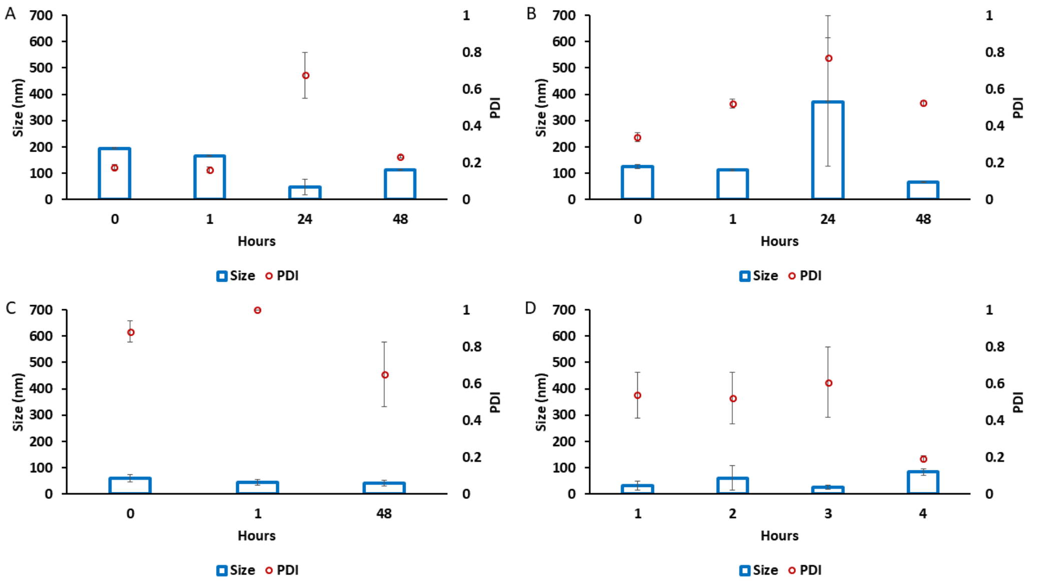


**Figure S7. Intensity weighted particle size and dispersity of Nano-BDNF B2.** The complexes were prepared by manual mixing of the solutions of BDNF and PSR-PLE in 10 mM HEPES, pH 7.4 at the $Z_{-/+}$ ratios of (A) 1, (B) 2, (C) 5, and (D) 10. The DLS measurements were carried out for up to 48 hours. Values are mean ± SEM.

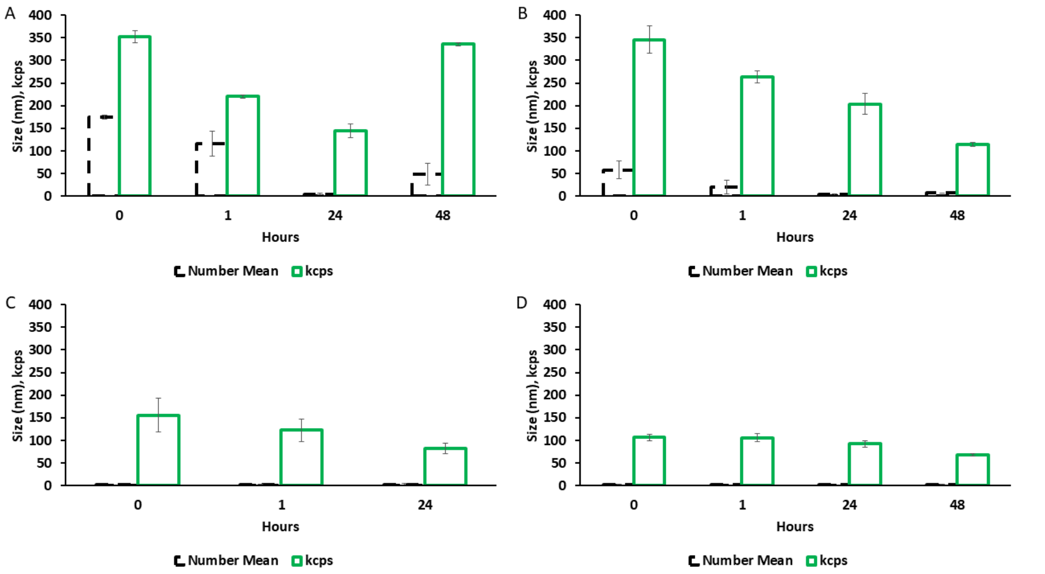


**Figure S8. Number weighted particle size and kilocount per second of Nano-BDNF B2 measurements in DLS of the same samples as shown Fig. S7.**

*Low ionic strength, minimal buffer for B2 (Fig. S7 and S8).*

Nano-BDNF B2 ($PMeOx_7$-$PPaOx_{29}$-$PMeOx_9$) formulated in 10 mM HEPES, pH 7.4 generally formed small particles with high PDI. $Z_{-/+}$=1 yielded the best results though there was a notable outlier at 24 hours. This high dispersity repeated in a second trial and may be indicative of a higher order process on a longer time scale. However, as the purpose was to identify the best candidates for optimization, we selected B2 in addition to others to move forward.

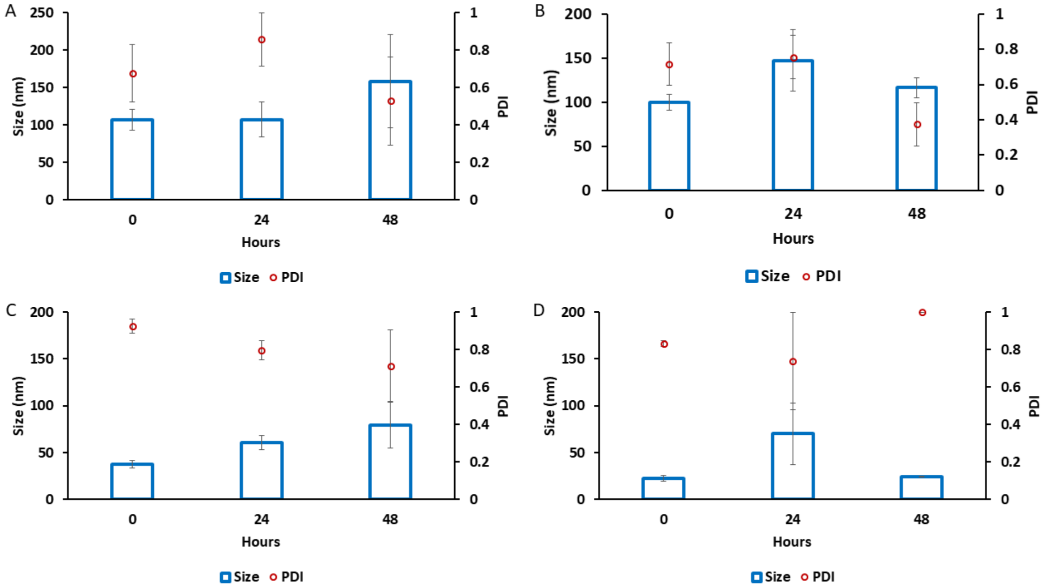


**Figure S9. Intensity weighted particle size and dispersity of Nano-BDNF B3.** The complexes were prepared by manual mixing of the solutions of BDNF and PSR-PLE in 10 mM HEPES, pH 7.4 at the $Z_{-/+}$ ratios of (A) 1, (B) 2, (C) 5, and (D) 10. The DLS measurements were carried out for up to 48 hours. Values are mean ± SEM.

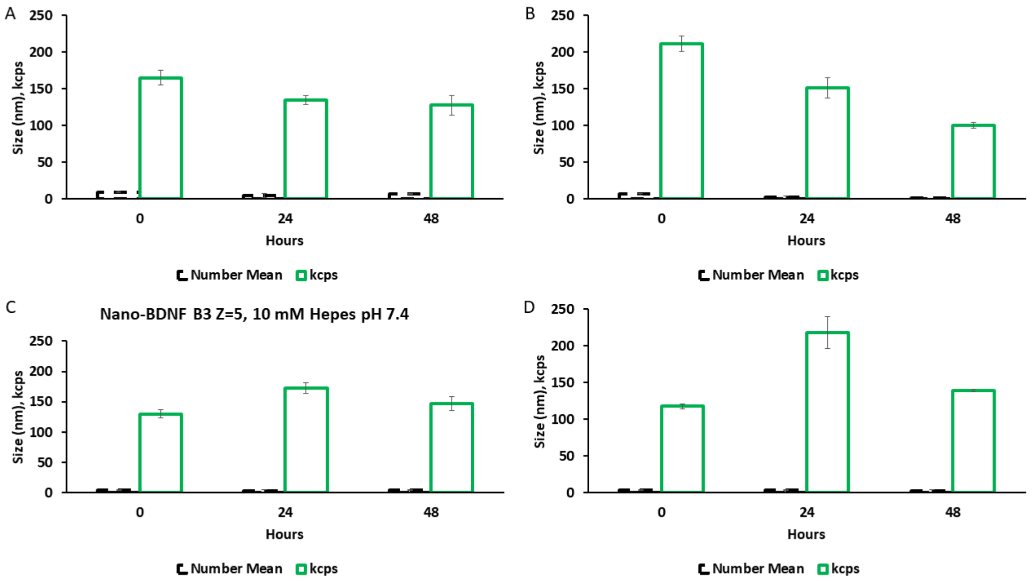


**Figure S10. Number weighted particle size and kilocount per second of Nano-BDNF B3 measurements in DLS of the same samples as shown Fig. S9.**

*Low ionic strength, minimal buffer for B3 (Fig. S9 and S10)*

Nano-BDNF B3 ($PMeOx_{33}$-$PPaOx_{10}$-$PMeOx_{31}$) formulated in 10 mM HEPES, pH 7.4 generally formed small particles with high PDI. The dispersity was high in all cases and we eliminated B3 from consideration.

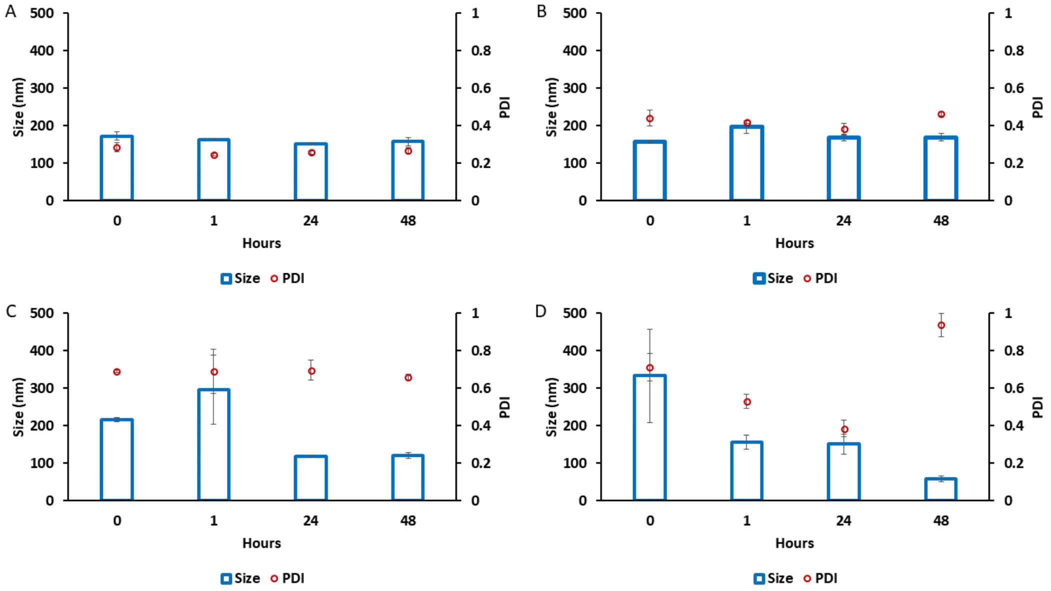


**Figure S11. Intensity weighted particle size and dispersity of Nano-BDNF B5.** The complexes were prepared by manual mixing of the solutions of BDNF and PSR-PLE in 10 mM HEPES, pH 7.4 at the $Z_{-/+}$ ratios of (A) 1, (B) 2, (C) 5, and (D) 10. The DLS measurements were carried out for up to 48 hours. Values are mean ± SEM.

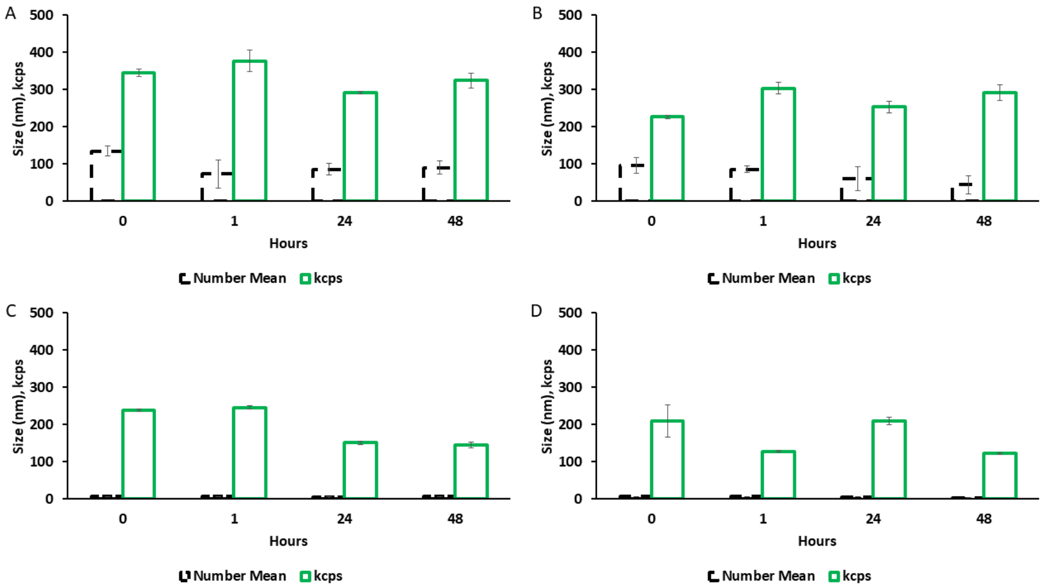


**Figure S12. Number weighted particle size and kilocount per second of Nano-BDNF B5 measurements in DLS of the same samples as shown Fig. S11.**

*Low ionic strength, minimal buffer for B5 (Fig. S11 and S12)*

Nano-BDNF B5 ($PMeOx_{38}$-$PPaOx_{27}$-$PMeOx_{38}$) formulated in 10 mM HEPES, pH 7.4 generally formed small particles with relatively low PDI at low $Z_{-/+}$ ratio. We continued optimization with B5.

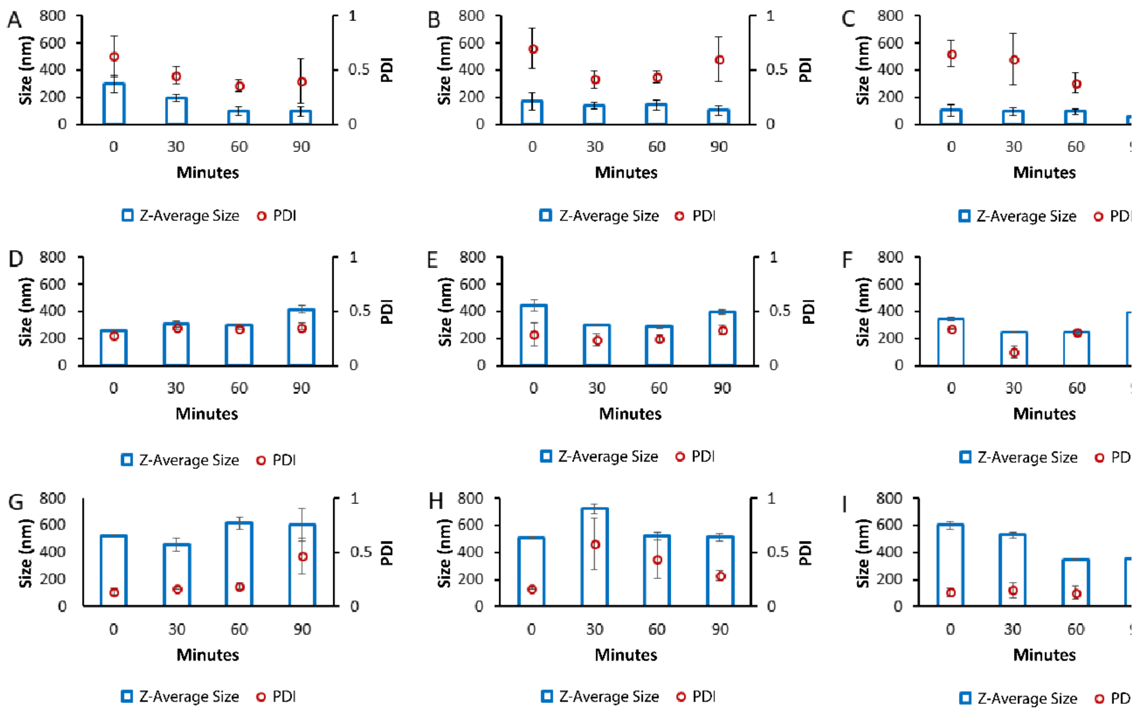


**Figure S13. Intensity weighted particle size and dispersity of Nano-BDNF in the presence of small-molecular mass electrolyte.** Nano-BDNF (A) B2 $Z_{-/+}$=1, (B) B2 $Z_{-/+}$=2, (C) B2 $Z_{-/+}$=5, (D) B5 $Z_{-/+}$=1, (E) B5 $Z_{-/+}$=2, (F) B5 $Z_{-/+}$=5, (G) PSR-PLE $Z_{-/+}$=1, (H) PSR-PLE $Z_{-/+}$=2, and (I) PSR-PLE $Z_{-/+}$=5 were prepared in 10 mM HEPES, 150 mM NaCl, pH 7.4 via manual mixing. The particles were measure for size and dispersity over 90 minutes using the Malvern Zetasizer DLS machine. Values are mean ± SEM.

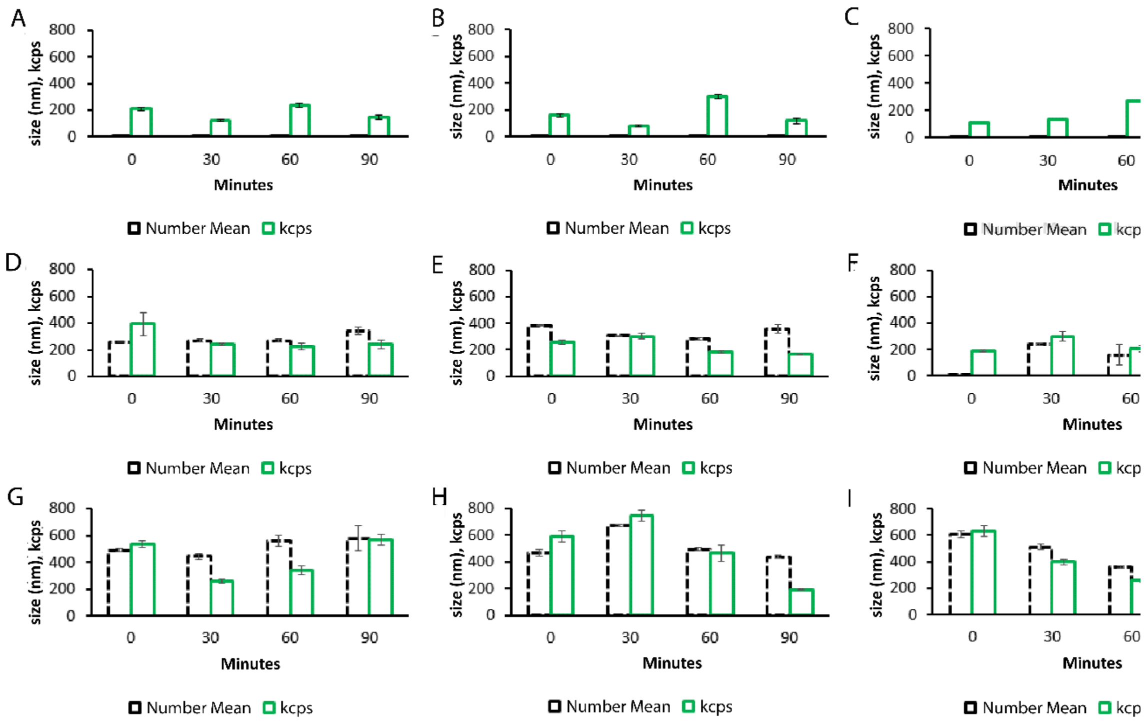


**Figure S14. Number weighted particle size and kilocount of Nano-BDNF in the presence of small-molecular mass electrolyte measured by DLS of the same samples as shown Fig. S13.** Nano-BDNF (A) B2 Z-/+=1, (B) B2 Z-/+=2, (C) B2 Z-/+=5, (D) B5 Z-/+=1, (E) B5 Z-/+=2, (F) B5 Z-/+=5, (G) PSR-PLE Z-/+=1, (H) PSR-PLE Z-/+=2, and (I) PSR-PLE Z-/+=5 were prepared in 10 mM HEPES, 150 mM NaCl, pH 7.4 via manual mixing. The particles were measure for size and dispersity over 90 minutes using the Malvern Zetasizer DLS machine. Values are mean ± SEM.

*High Ionic Strength buffer formulation of remaining candidate formulations (Figs. S13 and S14)*

We formulated our remaining candidate formulations, Nano-BDNF B2, B5, and PSR-PLE using 10 mM Hepes, 150 mM NaCl, pH 7.4. Higher ionic strength was expected to affect the formation of the PIC though the effects are difficult to predict. The effects of NaCl on PICs and other electrostatically driven aggregates can vary. Sometimes it can promote formation of small nanoparticles with low dispersity and other times it can prevent controlled association or association at all. In our case we observed that some conditions improved and others worsened.

Most notably, the higher $Z_{-/+}$ ratios of B5 and PSR-PLE (**Figure S13** and **S14**) were smaller and less disperse than in 10 mM HEPES, pH 7.4. It may be that the addition of NaCl allowed ion exchange and facilitated packing. B2 though had consistently high PDI though we continued work to observe the effect of microfluidic mixing. Notably, the mean number size for B2 was significantly lower indicating that the presence of salt promoted the presence of free polymer and BDNF.

*S1.4 Particle stability in isotonic solutions*

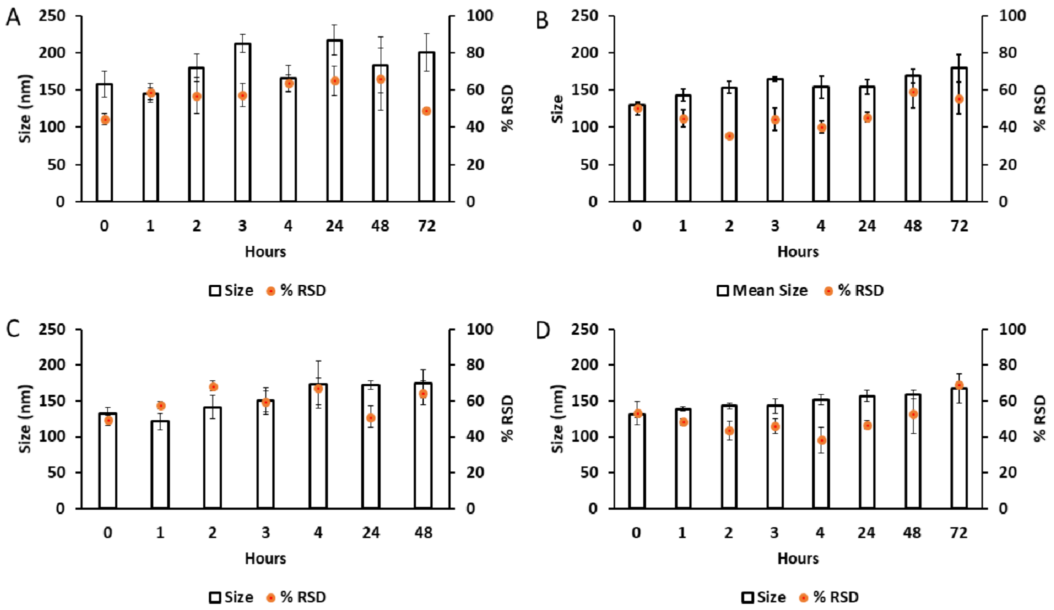


**Figure S15: Particle size and distribution of mixer formulated Nano-BDNF in (A,C) PBS and (B,D) LR solution over time.** (A,B) Nano-BDNF PSR-PLE $Z_{-/+}$=5 and (C,D) Nano-BDNF B5 $Z_{-/+}$=5 were analyzed at the indicated times using the Zetaview machine. Data is presented as mean ± SEM.

*S1.5 Titration of block copolymers*

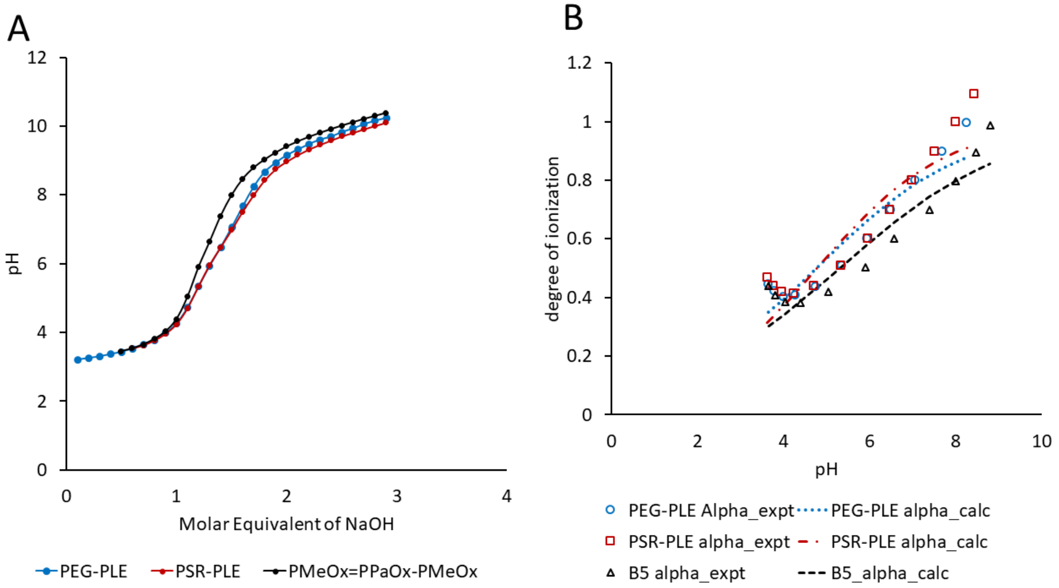


**Figure S16. Titration of Block Copolymers.** (A) The curves of pH titration using NaOH and (B) the degree of ionization for the three block copolymers. The degree of ionization was determined experimentally from the titration data or fitted to the experimental data as described below.

Table S1. The calculated effective $pK_a$ values and degrees of ionization at critical pH values[a]

| Block Copolymer | Effective $pK_a$ | Hill coefficient, *n* | Degree of Ionization, α | | |
|---|---|---|---|---|---|
| | | | **pH 5.0** | **pH 5.8** | **pH 7.4** |
| PEG-PLE | 4.75 | 0.25 | 0.54 | 0.65 | 0.82 |
| PSR-PLE | 4.78 | 0.29 | 0.54 | 0.66 | 0.85 |
| B5 | 5.29 | 0.22 | 0.46 | 0.56 | 0.74 |

a Effective pKa and α are defined here from the fitted modified Henderson Hasselbalch equation as described below.

.

Understanding the effects of pH on our polymers is critical in the characterization of our complex systems. We characterized the degree of ionization for each polymer and calculated the $pK_a$ values to better understand the effects of our perturbations of the system. We fitted the modified Henderson Hasselbalch equation using the least squares fit in Microsoft Excel to our degree of ionization data and thus obtained predicted values at pH 5.0, 5.8, and 7.4.

$$\alpha(pH) = \frac{1}{10^{n(pKa-pH)}+1}, \tag{S-1}$$

Where α is the degree of ionization, $pK_a$ is an effective $pK_a$ and n is the Hill coefficient[2,3].

After obtaining the titration curve, we identified a region of low ionization which would serve as an effective point zero for tracking the molar ratio of added base (γ) as compared to the concentration of ionizable units ($c_m$):

$$\gamma = \frac{[NaOH]}{c_m}, \tag{S-2}$$

The $c_m$ is the concentration of the carboxylic groups from the repeating units of the anionic block copolymers the numbers of which were previously determined by the $^1$H NMR spectra (B5) or provided by the manufacturer (PEG-PLE, PSR-PLE).

We used the relationship (S-3) whereby we can relate the degree of ionization (α) to (γ) as comes from charge neutralization:

$$\alpha = \gamma + \frac{[H^+]-[OH^-]}{c_m}, \tag{1}$$

This was then fitted using a least squares analysis using the relationship between pH and α (S-1).

*S1.6 Supplementary ITC Data*

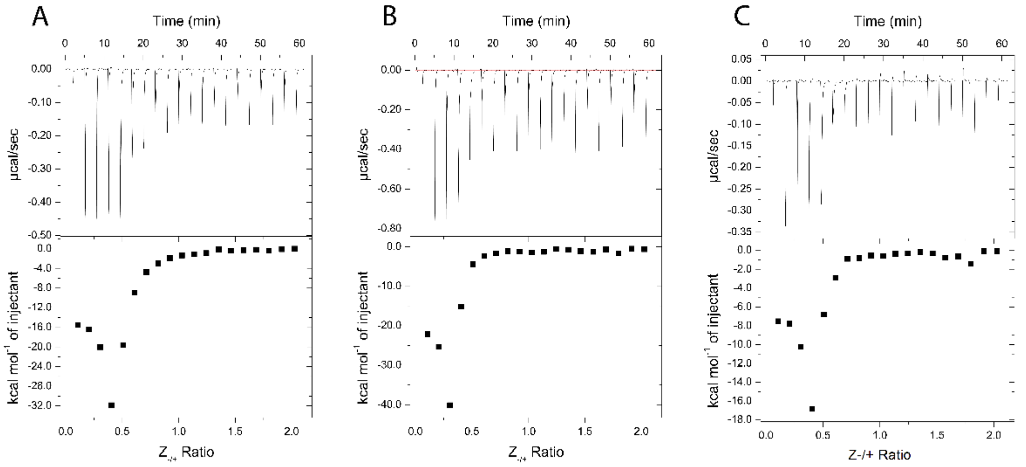


**Figure S17. ITC thermograms of Nano-BDNF formation in 10 mM sodium phosphate buffer, pH 5.8.** The holding cell was filled with 200 µL of 10 µM BDNF in at 25°C. (A) 88 µM PEG-PLE (B) 88 µM PSR-PLE, and (C) 162 µM B5 E were injected in 2 µL increments with a 3 min dwell time between each injection. Top: time dependence of the heat supplied to the sample cell for each injection of the block copolymer into the BDNF solution. Bottom - integrated ITC curves for the block copolymer binding with BDNF as a function of the charge ratio $Z_{-/+}$.

**Table S2.** Critical $Z_{-/+}$ values of in the analysis of ITC thermograms.

| Block Copolymer | $Z_{-/+}$ of the curve minimum | | | $Z_{-/+}$ of the curve saturation | | |
|---|---|---|---|---|---|---|
| | **pH 7.4** | **pH 5.8** | **pH 5.0** | **pH 7.4** | **pH 5.8** | **pH 5.0** |
| PEG-PLE | 0.4 | 0.4 | 0.6 | 0.9 | 1.0 | 2.2 |
| PSR-PLE | 0.4 | 0.3 | 0.6 | 0.6 | 0.6 | 2.0 |
| B5 | 0.4 | 0.4 | 1.0 | 0.9 | 0.7 | 2.2 |

1. Jiang, Y.; Fay, J.M.; Poon, C.D.; Vinod, N.; Zhao, Y.; Bullock, K.; Qin, S.; Manickam, D.S.; Yi, X.; Banks, W.A.; et al. Nanoformulation of Brain-Derived Neurotrophic Factor with Target Receptor-Triggered-Release in the Central Nervous System. *Adv. Funct. Mater.* **2018**, *28*, 1–11, doi:10.1002/adfm.201703982.

2. Borukhov, I.; Andelman, D.; Borrega, R.; Cloitre, M.; Leibler, L.; Orland, H. Polyelectrolyte Titration: Theory and Experiment. *J. Phys. Chem. B* **2000**, *104*, 11027–11034, doi:10.1021/jp001892s.

3. Bodnarchuk, M.S.; Doncom, K.E.B.; Wright, D.B.; Heyes, D.M.; Dini, D.; O'Reilly, R.K. Polyelectrolyte PKa from Experiment and Molecular Dynamics Simulation. *RSC Adv.* **2017**, *7*, 20007–20014, doi:10.1039/c6ra27785c.
